# High-density Lipoprotein Inspired Lipid Nanoparticles for Systemic RNA delivery to the Brain

**DOI:** 10.64898/2026.05.29.728695

**Authors:** Zunkai Xu, Kiruba Shalin, Vy Do, Alexander Yan, Jingjing Gao

## Abstract

Systemic delivery of nucleic acids to the central nervous system (CNS) remains fundamentally limited by the blood-brain barrier (BBB) and the lack of cell-type specificity. Inspired by endogenous high-density lipoproteins (HDLs) that naturally traverse the BBB, we engineered a library of 112 HDL-inspired lipid nanoparticles (HLNPs) to deliberately program protein corona composition and direct nanoparticle trafficking across the BBB. High-throughput *in vivo* screening identified HLNPs with robust brain accumulation and preferential enrichment within distinct neural cell populations. Quantitative proteomic analysis revealed that elevated apolipoprotein A-I/apolipoprotein E and vitronectin/apolipoprotein E ratios correlate with enhanced BBB transport. We further demonstrated that neuron-targeted HLNPs delivering PTEN siRNA produced functional recovery and lesion reduction in a murine traumatic brain injury model, while microglia-targeted HLNPs carrying interleukin-10 mRNA potently suppressed neuroinflammation. Together, these findings establish HLNPs as a versatile RNA delivery platform that leverages protein corona programming for enhanced brain delivery and cell-type-selective targeting.

## Main

RNA-based therapeutics have emerged as a transformative class of medicines, offering sequence-level precision in modulating gene expression through mechanisms inaccessible to conventional small molecules or protein biologics^1,2^. Key modalities include small interfering RNAs (siRNAs) for targeted gene silencing, messenger RNAs (mRNAs) for transient therapeutic protein expression, and microRNA mimics or antisense oligonucleotides (ASOs) for broad regulation of disease-associated gene networks^3,4^. The CNS represents one of the most compelling frontiers for RNA medicine: neurological diseases including Alzheimer’s disease, Parkinson’s disease, ALS, traumatic brain injury, and neuroinflammatory disorders collectively affect over one billion people worldwide and impose an enormous unmet clinical burden. Many of these conditions are driven by well-characterized gain-of-function mutations, dysregulated gene networks, or pathological inflammatory signaling that are ideally suited to siRNA-mediated silencing or mRNA-based protein replacement^5,6^.

Despite this extraordinary potential, the clinical translation of RNA therapeutics for CNS diseases faces formidable and interconnected delivery challenges. RNA molecules are inherently unstable in biological fluids, rapidly degraded by serum nucleases following systemic administration, and their large size and polyanionic charge prevent passive membrane crossing^7^. Most critically, the brain is protected by the blood-brain barrier (BBB), a highly specialized neurovascular interface of tight junction-coupled endothelial cells, pericytes, and astrocytic endfeets that excludes over 98% of systemically administered therapeutics, including virtually all nucleic acids, from reaching the brain parenchyma at meaningful concentrations^8,9^. Beyond the BBB, the brain comprises functionally distinct cell populations including neurons, microglia, astrocytes, and oligodendrocytes, each playing specific and often opposing roles in CNS disease^10^. Non-selective RNA delivery across all brain cell types dilutes therapeutic efficacy and risks off-target gene silencing or inflammatory activation in healthy populations^11^. Together, nuclease susceptibility, BBB exclusion, and lack of cell-type specificity define the central translational bottleneck for CNS RNA medicine.

Lipid nanoparticles (LNPs) represent the most clinically advanced RNA delivery platform, validated by the approval of patisiran for hepatic RNA interference and the global deployment of mRNA-LNP COVID-19 vaccines^12,13^. However, a fundamental limitation of current LNPs is their hepatotropic bias: systemic administration drives rapid ApoE adsorption from plasma, which mediates preferential liver accumulation through LDL receptor engagement and severely restricts extrahepatic utility^14,15^. Recent efforts to redirect LNPs toward the brain include adding BBB-crossing lipids for CNS mRNA delivery^16^ and furan-derived LNPs exploiting meningeal lymphatic vessels as an alternative brain entry route^17^, yet broader strategies using surface-conjugated targeting ligands or receptor-mediated transcytosis remain hampered by receptor saturation, lysosomal cargo degradation, and poor clinical translatability^18^. Critically, none of these approaches has achieved programmable, cell-type-selective RNA delivery within the brain parenchyma, and no systemically administered RNA therapeutic has received regulatory approval for any CNS indication, underscoring a fundamental unresolved gap in the field^19^.

A critical but underexplored determinant of LNP biodistribution is the protein corona, the dynamic layer of plasma proteins adsorbed onto nanoparticle surfaces upon entry into the bloodstream, which fundamentally dictates receptor interactions, cellular uptake pathways, and organ-level distribution^20–22^. Although corona composition is highly sensitive to nanoparticle surface chemistry and lipid content, its rational exploitation to reprogram LNP trafficking toward the BBB remains largely uninvestigated^23,24^. Intriguingly, the bloodstream harbors a naturally occurring nanoparticle that traverses the BBB: high-density lipoprotein (HDL)^25^. HDL is a dynamic nanoscale complex of phospholipids and apolipoproteins, mainly apolipoprotein A-I (ApoA-I), that engages SR-B1 receptors highly expressed on brain endothelial cells, enabling BBB crossing through receptor-mediated transcytosis^26,27^. We therefore reasoned that incorporating HDL-derived phospholipids into LNP formulations could bias protein corona composition toward BBB-relevant apolipoproteins, while attenuating hepatotropic ApoE adsorption, thereby programming nanoparticle trafficking across the BBB.

To implement this strategy, we analyzed the lipid composition of native HDL and selected seven HDL-mimetic lipids, four cholesterol derivatives, and four clinically validated ionizable lipids to construct a 112-member HLNP library. High-throughput *in vivo* screening identified lead HLNPs with markedly enhanced brain accumulation compared with the clinically established Patisiran-based liver-targeting LNP formulation (hereafter “Pati”), and revealed distinct formulations with preferential enrichment in specific neural cell populations. Quantitative proteomic profiling identified ApoA-I, vitronectin, and ApoE as consistently abundant corona proteins, with HLNPs exhibiting the highest brain accumulation showing elevated ApoA-I/ApoE and vitronectin/ApoE ratios, corroborating the HDL-mimetic design principle and implicating these signatures as predictive biomarkers of BBB transport efficiency (**Fig. 1a**). Building on these insights, we characterized cell-type-specific RNA delivery profiles across multiple brain cell populations and prioritized neuron- and microglia-targeting HLNPs for therapeutic validation in murine models of traumatic brain injury (TBI) and LPS-induced neuroinflammation, respectively. Collectively, this work establishes HDL-inspired lipid nanoparticles as a clinically translatable platform for targeted RNA delivery to the CNS, with broad applicability across neurological disorders driven by distinct cell-type-specific pathologies.

**Fig. 1:**
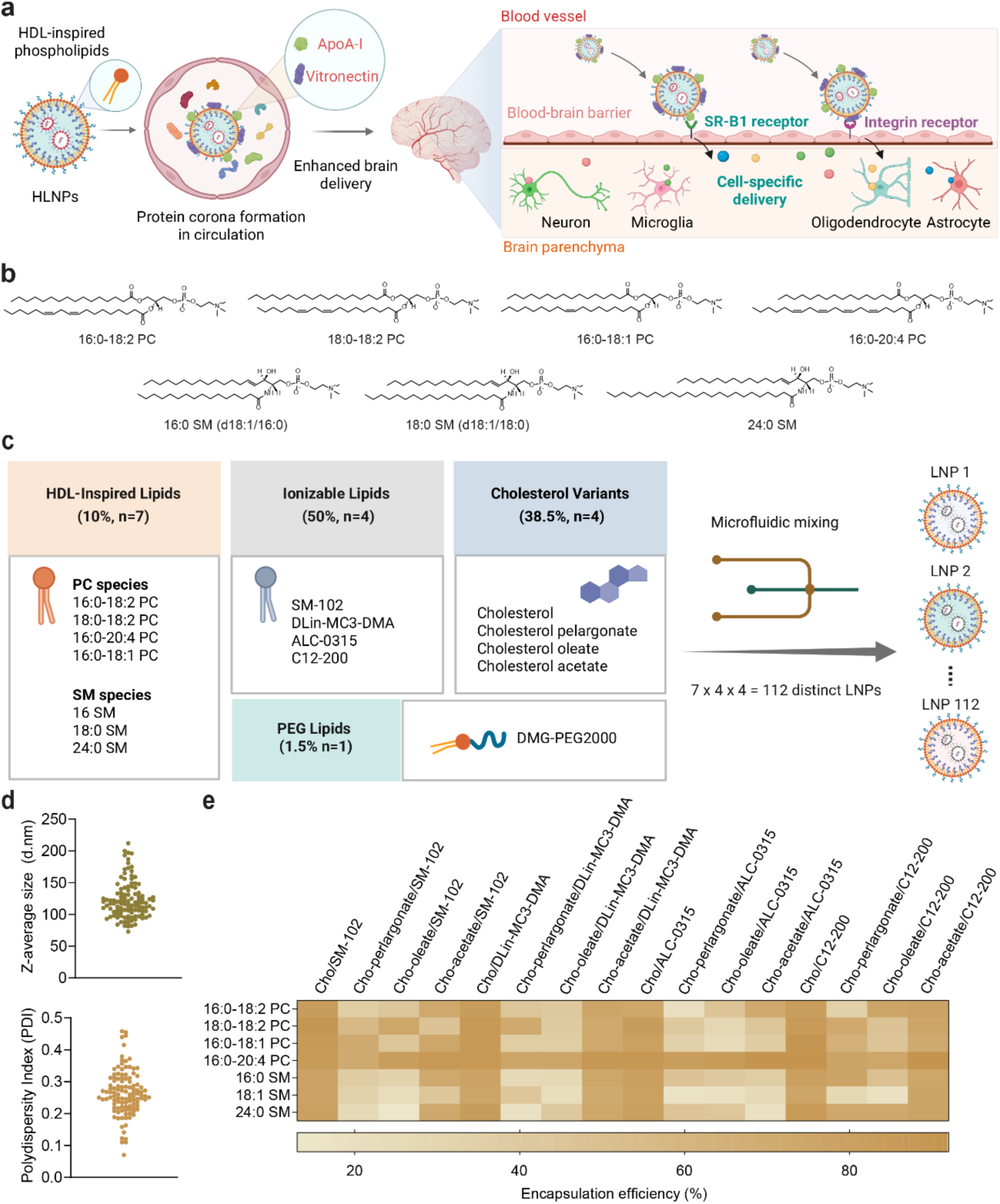
Construction and physicochemical characterization of an HLNP library. **a,** Schematic illustration of the design rationale and proposed delivery mechanism of HLNPs. HLNPs formed protein coronas enriched in apolipoproteins and vitronectin in circulation, facilitating receptor-mediated docking at the BBB via SR-BI and integrin receptors, followed by transcytosis and subsequent profiling of distribution across different brain cell types in the parenchyma. **b**, Chemical structures of selected HDL phospholipids used to construct the HLNP library. **c**, Combinatorial design of the HLNP library. A single lipid from each category: HDL-mimetic phospholipids, ionizable lipids, cholesterol variants and a DMG-PEG2000, were combined *via* microfluidic mixing to generate 112 chemically distinct formulations. **d**, Physicochemical characterization of the HLNP library, showing hydrodynamic diameter and PDI distributions across formulations. **e,** Heatmap of encapsulation efficiency across HLNP formulations, organized by phospholipid identity (rows) and ionizable lipid/cholesterol combinations (columns), highlighting formulation-dependent variations in cargo loading.

### Rational design and construction of HLNPs for brain delivery

Guided by the lipid compositional profile of native HDL^28^, we constructed a library of HLNPs by systematically incorporating lipids from three functionally distinct components. First, to recapitulate the phospholipid surface of HDL, we selected four representative phosphatidylcholine (PC) species (16:0-18:2 PC, 18:0-18:2 PC, 16:0-20:4 PC and 16:0-18:1 PC) and three sphingomyelin (SM) species (16:0 SM, 18:0 SM and 24:0 SM) , which together span the dominant phospholipid classes found in native HDL and enable modulation of membrane fluidity, lipid packing, and nanoparticle surface organization to regulate protein corona composition (**Fig. 1b**). Second, to modulate membrane rigidity and lipoprotein-like surface architecture, we incorporated four cholesterol analogues (cholesterol, cholesterol pelargonate, cholesterol oleate, and cholesterol acetate; **Supplementary Fig. 1**), which vary in acyl chain length and ester polarity and are known to differentially influence sterol packing, membrane order, and nucleic acid encapsulation efficiency^29–31^. Third, to enable endosomal escape and cytoplasmic RNA delivery, we included four clinically validated ionizable lipids (SM-102, DLin-MC3-DMA, ALC-0315, and C12-200; **Supplementary Fig. 2**), whose distinct pKa values and lipid geometries govern pH-dependent charge switching, membrane fusion activity, and intracellular delivery efficiency^20^. Using microfluidic mixing, we implemented an orthogonal combinatorial design in which each formulation comprised one HDL-inspired phospholipid (PC or SM), one cholesterol analogue, one ionizable lipid and DMG-PEG2000, generating a library of 112 HLNP formulations (**Fig. 1c**).

All 112 HLNP formulations were systematically characterized for particle size, polydispersity index (PDI), and encapsulation efficiency (**Supplementary Table 1**). Across the library, HLNPs exhibited uniform nanoscale diameters of approximately 100-200 nm with low polydispersity (PDI < 0.35; **Fig. 1d**). Encapsulation efficiency, however, varied substantially across formulations and was predominantly governed by the cholesterol component rather than the HDL-derived phospholipid identity (**Fig. 1e**). Formulations incorporating cholesterol or cholesterol acetate achieved higher encapsulation efficiencies (approximately 70-90%) compared with those containing cholesterol oleate or cholesterol pelargonate. This pattern is consistent with the known influence of sterol structure on lipid packing and membrane organization: the shorter, less polar acyl chains of cholesterol acetate relative to the longer, more hydrophobic chains of cholesterol oleate and pelargonate likely produce tighter lipid packing and a more ordered bilayer, creating a more enclosed hydrophobic core that retains nucleic acid cargo more efficiently^32–34^. In contrast, variation in HDL-derived phospholipid species had minimal influence on encapsulation, suggesting that the phospholipid component primarily shapes surface properties rather than cargo retention. Based on these characterization data, formulations meeting predefined thresholds for particle size (100-200 nm) and PDI (< 0.35) were further ranked by encapsulation efficiency, and the top 66 candidates were advanced for *in vivo* screening.

### High-throughput *in vivo* screening of HLNPs for brain delivery

To enable high-throughput *in vivo* screening, each HLNP formulation was encapsulated with a unique barcode DNA (b-DNA) for sequencing-based biodistribution analysis^35^. A total of 66 HLNPs were grouped into three pools and intravenously administered to C57BL/6 mice, enabling unbiased head-to-head comparison of brain delivery efficiency across formulations (**Fig. 2a**). Heatmap visualization of b-DNA abundance revealed significant formulation-dependent heterogeneity across various organs (**Fig. 2b**) with the ranking of individual formulations provided in **Supplementary Figs. 3-7**.

**Fig. 2:**
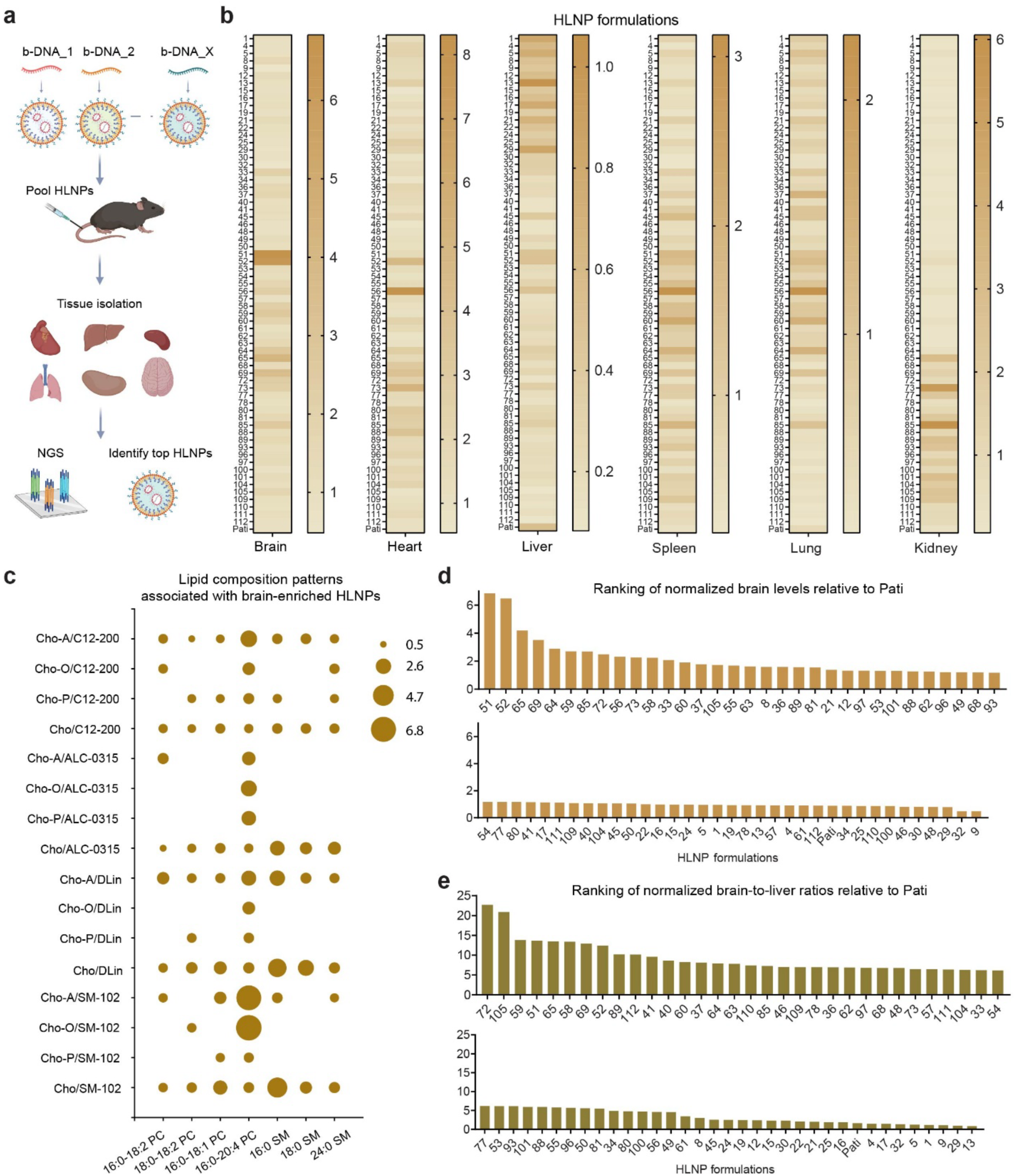
Multiplexed *in vivo* screening identifies brain-enriched HLNPs. **a**, Schematic illustration of the *in vivo* screening workflow for pooled barcoded HLNPs. Distinct HLNP formulations, each encapsulating a unique b-DNA, were divided into three pooled batches and intravenously administered to mice. Four hours after intravenous administration, major organs were collected, b-DNA were recovered and amplified, and deep sequencing was performed to quantify formulation-specific biodistribution. **b**, Heatmap showing the relative abundance of 66 HLNP formulations across major organs, including brain, heart, liver, spleen, lung and kidney. Each row represents an individual HLNP formulation, and color gradients indicate normalized barcode abundance recovered from each tissue. Values were independently normalized within each tissue, and thus are not directly comparable across organs. **c**, Bubble plot summarizing lipid composition features of HLNP formulations identified from the *in vivo* screen, highlighting phospholipid-dependent trends associated with brain enrichment (Cho, cholesterol; Cho-P, cholesterol pelargonate; Cho-O, cholesterol oleate; Cho-A, cholesterol acetate). **d,** Ranking of HLNP formulations according to brain-associated b-DNA abundance that are normalized to benchmark Patisiran formulation. **e**, Ranking of HLNP formulations according to brain-to-liver enrichment ratios of b-DNA normalized to benchmark Patisiran formulation.

Within the brain, several formulations, notably HLNPs 51, 52, 65 and 69, displayed markedly elevated barcode signals relative to the rest of the library. In contrast, a subset of formulations, including HLNPs 13, 29, 1 and 17, exhibited preferential accumulation in the liver. Distinct enrichment patterns were also observed across other organs. To further elucidate structure-function relationships driving brain enrichment, we examined the lipid composition of representative HLNP formulations. Bubble plot analysis revealed that brain-enriched HLNPs were associated with certain specific phospholipid species, particularly unsaturated 16:0-20:4 PC and 16:0 SM. In contrast, trends for other lipid components, including cholesterol derivatives and ionizable lipids, were less pronounced (**Fig. 2c**). These findings suggest that phospholipid identity is a primary determinant of brain delivery, potentially through modulation of protein corona composition and subsequent interactions with BBB transport pathways. Mechanistically, phospholipids bearing polyunsaturated acyl chains may enhance membrane fluidity and alter lipid packing, whereas sphingomyelin species can promote the formation of cholesterol-rich ordered domains, thereby reshaping nanoparticle surface organization and protein adsorption profiles^36^.

To identify lead candidates for brain delivery, HLNPs were ranked based on absolute brain abundance and brain-to-liver enrichment ratios, the latter of which accounts for the typical hepatic sequestration of systemically administered lipid nanoparticles. Brain ranking analysis revealed that several formulations, including HLNPs 51, 52, 65 and 69, exhibited the highest brain-associated barcode signals and clearly separated from the remainder of the library (**Fig. 2d**). Notably, HLNP 51 and HLNP 52 displayed brain signals approximately 14-fold and 13.5-fold higher, respectively, than the lowest-ranked formulation (HLNP 9), indicating a substantial dynamic range in brain accumulation across the HLNP library. Ranking based on brain-to-liver enrichment ratios further identified HLNPs 72, 105 and 59 as the top candidates, followed by HLNP 51, 65 and 58 (**Fig. 2e**). In this analysis, HLNP 72 and HLNP 105 exhibited brain-to-liver ratios approximately 26-fold and 24-fold higher, respectively, than the lowest-ranked formulation (HLNP 13), highlighting substantial variability in brain selectivity across the screened nanoparticles. In contrast, the clinically validated liver-targeting LNP formulation Pati exhibited minimal brain enrichment in this screening dataset, consistent with its strong hepatic tropism^37^.

### Top-ranked HLNPs mediate functional mRNA expression and siRNA-mediated gene silencing

To assess functional delivery, we evaluated the capabilities of our top-ranked HLNPs to deliver a luciferase mRNA reporter across neuronal, microglial and endothelial cell lines (**Fig. 3a–d**). N2a, BV2 and bEnd.3 cells were treated with individual formulations, and luciferase expression was quantified after 24 h. Several HLNPs mediated robust luciferase expression across cell types, with some exhibiting comparable or superior transfection efficiency relative to the Pati control. Interestingly, a discrepancy between *in vitro* activity and *in vivo* biodistribution was noted. Specifically, HLNPs 9 and 13, which showed low brain accumulation in previous *in vivo* screens, elicited relatively higher levels of luciferase activity *in vitro.* Successful delivery and expression of mCherry mRNA in N2a and BV2 cells also validated the functional capabilities of these HLNPs (**Fig. 3e**). In addition to mRNA delivery, we also assessed the capacity of these HLNPs for siRNA-mediated gene silencing using *Ppib*-targeting siRNA. Most HLNPs achieved efficient *Ppib* knockdown in N2a cells (**Fig. 3f,g**), indicating robust intracellular delivery and RNA interference activity. Together, these results demonstrate that top-performing HLNPs enable functional mRNA expression and siRNA-mediated gene silencing.

**Fig. 3:**
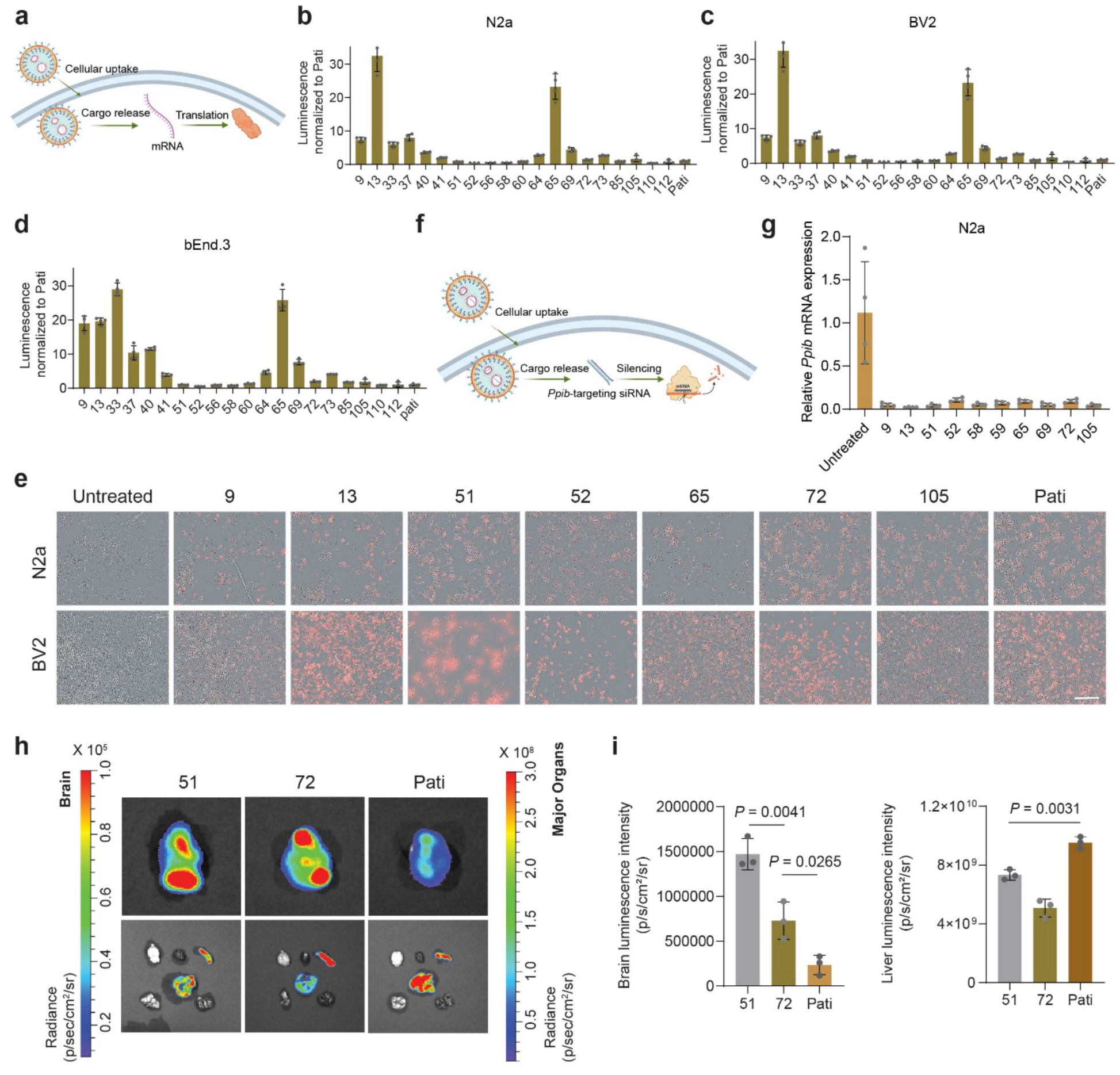
Functional evaluation of top HLNP-mediated mRNA expression and siRNA-mediated gene silencing. **a**, Schematic illustration of HLNP-mediated mRNA delivery and translation. **b-d**, Luciferase activity of HLNP treated N2a neuronal cells (b, Samples 56 and 110 had *n* = 3; all other groups had *n* = 4), BV2 microglial cells (c, Samples 52, 56, 58, 60 and 69 had *n* = 3; all other groups had *n* = 4) and bEnd.3 brain endothelial cells (d, *n* = 4). **e**, Representative fluorescence images of N2a and BV2 cells following delivery of mCherry mRNA by selected HLNP formulations. Blank indicates untreated cells. Scale bar, 100 μm. **f**, Schematic illustration of HLNP-mediated siRNA delivery. **g**, Relative *Ppib* mRNA levels in N2a cells following siRNA delivery, quantified by RT-qPCR and normalized to housekeeping gene *Gapdh* (*n* = 4). **h,** Representative *ex vivo* bioluminescence images of isolated brains collected from mice following intravenous administration of luciferase mRNA-loaded LNPs (51, 72, and Pati) (top). Corresponding *ex vivo* images of major organs (bottom) show distinct biodistribution patterns among the formulations. **i**, Quantification of normalized luminescence intensity in the brain and liver based on the ex vivo imaging shown in panel h (*n* = 3).

To further validate functional delivery *in vivo*, selected HLNPs 51 (top 1 based on brain abundance) and 72 (top 1 based on brain/liver ratio enrichment) and the liver-targeting control Pati were evaluated individually *in vivo* using a luciferase mRNA reporter (**Fig. 3h**). Ex vivo bioluminescence imaging of isolated brains and major organs following luciferin administration revealed markedly stronger brain-associated signals in mice treated with HLNPs 51 and 72, whereas the Pati formulation produced minimal cranial luminescence. Quantitative analysis demonstrated that HLNP 51 and HLNP 72 generated 6.3-fold and 3.4-fold higher brain luminescence intensities, respectively, than Pati. Notably, while HLNP 51 produced strong signals in both brain and liver, HLNP 72 exhibited comparably high brain expression but substantially reduced liver-associated luminescence, indicating improved brain selectivity. In contrast, the Pati formulation generated dominant liver signals with minimal brain expression (**Fig. 3i**). These results are consistent with the sequencing-based ranking from the pooled screening, confirming that leading HLNP formulations enable enhanced brain accumulation and functional mRNA delivery following systemic administration.

### Protein corona composition influences HLNP brain delivery

The pronounced differences in brain accumulation across HLNP formulations prompted us to investigate whether these biodistribution patterns arise from intrinsic nanoparticle properties or factors acquired dynamically in the bloodstream. To distinguish between these possibilities, we administered the same pooled HLNP library by two routes that differ specifically in plasma exposure: intravenous injection, which subjects nanoparticles to the full circulating plasma environment, and intrathecal injection, which delivers nanoparticles directly into the cerebrospinal fluid bypassing systemic circulation. These two routes produced markedly different organ distribution patterns (**Fig. 4a,b and Supplementary Fig. 8**), which strongly suggest that bloodstream-acquired protein coronas as a critical determinant of brain biodistribution *in vivo*. To characterize the protein corona formed under physiological conditions, selected HLNPs were formulated to encapsulate carboxylated iron oxide nanoparticles, enabling magnetic isolation of nanoparticle-protein complexes following incubation with mouse plasma (**Fig. 4c,d**). The recovered nanoparticle-associated proteins were then characterized by quantitative proteomic analysis^38^. For this analysis, we selected the top three HLNPs ranked by absolute brain accumulation (HLNP 51, 52 and 65), the top two formulations based on brain-to-liver ratio (HLNP 72 and 105), two bottom-ranked variants (HLNP 9 and 13), and Pati, a clinically validated liver-targeting LNP formulation, as comparators.

**Fig. 4:**
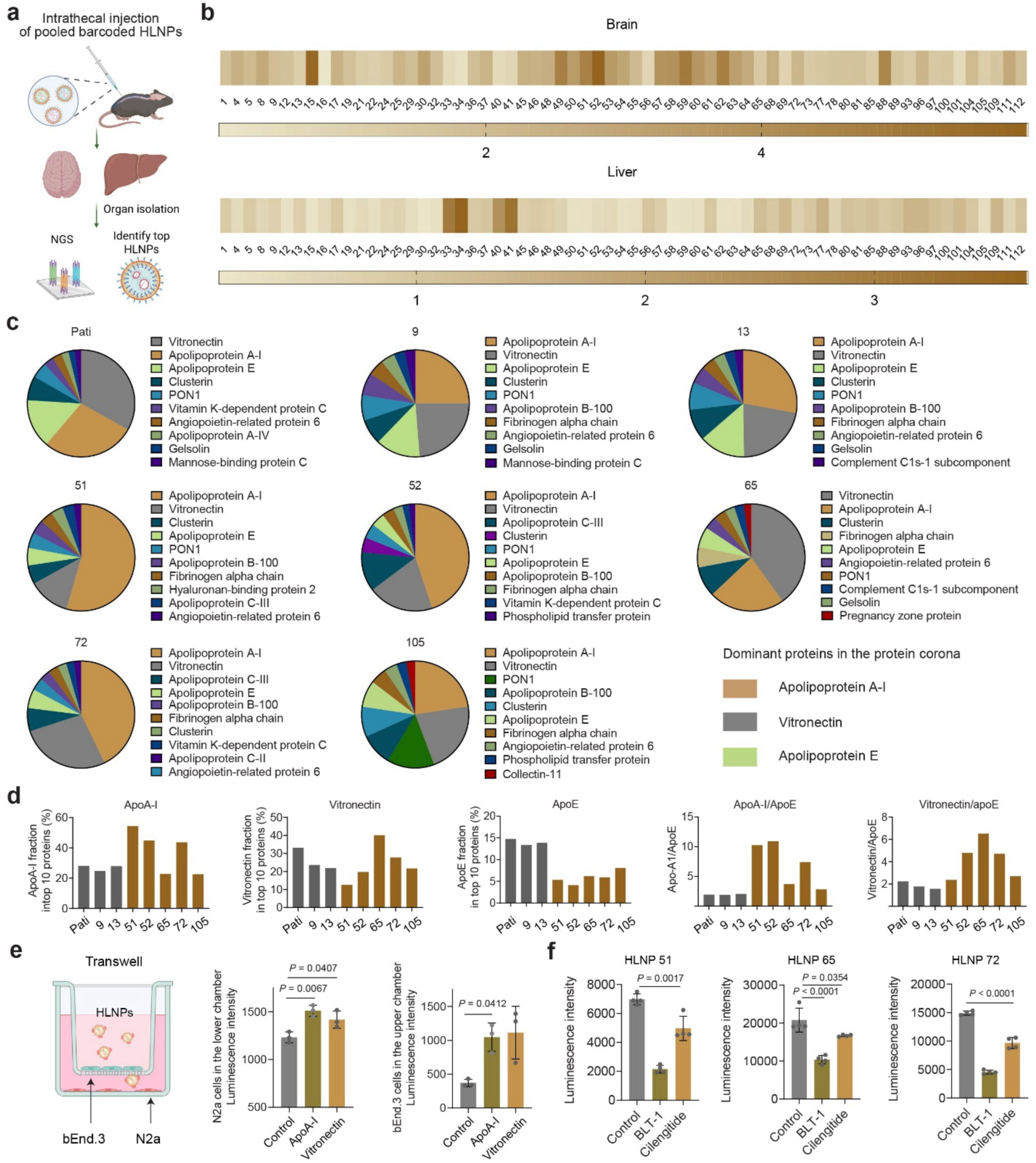
Protein corona–mediated mechanisms underlying HLNP brain delivery. **a,** Schematic illustration of intrathecal administration of pooled HLNPs. **b,** Heatmap of HLNP accumulation in the brain and liver following intrathecal injection, revealing distribution patterns distinct from systemic delivery. Values are independently normalized within each tissue. **c,** Proteomic analysis of the protein corona associated with representative HLNP formulations exhibiting distinct *in vivo* targeting profiles. Pie charts show the relative abundance of major adsorbed serum proteins, highlighting differential enrichment of ApoA-I, vitronectin and ApoE. **d,** Quantification of ApoA-I, vitronectin and ApoE abundance in the protein corona of selected HLNPs, along with corresponding ApoA-I/ApoE and VTN/ApoE ratios. Top-performing HLNPs show higher ratios compared with lower-performing formulations and the Pati control. **e,** Schematic illustration of HLNPs pre-coated with ApoA-I or vitronectin and evaluated in a transwell-based bEnd.3–N2a co-culture model. Luciferase expression was quantified following treatment with protein-precoated HLNP 51 (*n* = 4). **f,** Luciferase expression following treatment with representative HLNPs (51, 65 and 72) in the absence or presence of pharmacological inhibitors, including BLT-1 (SR-B1 inhibitor, 10 μM) and cilengitide (integrin αvβ3/αvβ5 antagonist, 5 μM) (*n* = 4).

The lowest brain-ranked (HLNP 9 and HLNP 13), and the liver-tropic Pati formulation exhibited markedly higher ApoE enrichment compared with other HLNPs, indicating a strongly ApoE-dominant corona profile. In contrast, several of the top-performing brain-directed HLNPs, including HLNP 51 and HLNP 52, displayed elevated ApoA-I levels and a substantially increased ApoA-I/ApoE ratio relative to the lower-performing formulations. In addition, HLNP 65, 72 and 105 showed pronounced enrichment of vitronectin compared with HLNP 9, 13 and Pati. Notably, although HLNP 72 and HLNP 105 did not exhibit the highest absolute levels of ApoA-I, they nevertheless showed markedly elevated ApoA-I/ApoE and vitronectin/ApoE ratios, a trend that was also observed in other top-performing formulations, including HLNP 51, 52, and 65. These results suggest that the balance among these three corona proteins, rather than the absolute abundance of any single component, is the key correlate of brain delivery efficiency.

These proteomic signatures are mechanistically consistent with known receptor biology at the BBB. ApoA-I, the principal structural apolipoprotein of HDL, engages scavenger receptor class B type I (SR-BI), which is highly expressed on brain endothelial cells and mediates surface-selective lipid exchange rather than classical endocytosis^25^. This SR-BI-mediated surface docking may facilitate sustained endothelial association and transcytosis initiation. ApoE, by contrast, engages LDL receptor family members, consistent with its association with hepatic sequestration rather than brain delivery^39^. Vitronectin, a plasma glycoprotein that bridges integrin receptors expressed at the neurovascular interface, may further stabilize nanoparticle docking at the luminal endothelial surface through integrin αvβ3/αvβ5 engagement^40^. Together, these observations suggest that ApoA-I and vitronectin enrichment cooperatively promotes productive BBB interactions, while ApoE dominance redirects nanoparticles toward hepatic clearance.

To assess the functional contribution of individual protein corona component, HLNP 51 encapsulating Cy5-labelled ssDNA was pre-incubated with recombinant ApoA-I or vitronectin prior to incubate with cells. Both proteins increased membrane association at 4 °C (**Supplementary Figs. 9,10**). In a transwell-based in vitro BBB model, pre-coating HLNP 51 with either ApoA-I or vitronectin enhanced luciferase expression in both endothelial and neuronal cells (**Fig. 4e**), indicating these corona proteins improved cellular interaction and uptake.

To probe uptake mechanisms, we next evaluated pharmacological inhibitors targeting candidate receptor pathways. Inhibition of SR-B1 with BLT-1 or blockade of integrins αvβ3/αvβ5 with cilengitide both reduced luciferase expression across representative HLNPs (51, 65 and 72) (**Fig. 4f**), implicating these receptors in corona-mediated HLNP uptake. Additional inhibition on endocytic pathways indicated that clathrin-mediated, cholesterol-dependent and dynamin-dependent processes contribute most substantially to HLNP internalization, whereas caveolae-mediated uptake and macropinocytosis play comparatively minor roles (**Supplementary Fig. 11**).

Collectively, these results establish a corona-dependent mechanism for HLNP brain delivery in which HDL-derived phospholipid composition biases protein adsorption toward an ApoA-I and vitronectin-enriched corona with reduced ApoE content, promoting SR-BI and integrin-mediated engagement at the brain endothelium and facilitating nanoparticle transcytosis into the brain parenchyma.

### *In vivo* screening of barcoded HLNPs defines cell-type–specific distribution profiles of top HLNPs

To define the cell-type-specific distribution of HLNPs within the brain, we extended the *in vivo* barcoded HLNP screening to a targeted 27 formulations including the top 18 formulations from brain accumulation and another top 9 formulations based brain-to-liver enrichment rankings from the initial *in vivo* screen. Following intravenous administration of pooled, barcoded HLNPs, brains were harvested, dissociated into single-cell suspensions, and five major neural and neurovascular cell populations were isolated by magnetic-activated cell sorting: neurons, microglia, astrocytes, oligodendrocytes, and cerebrovascular endothelial cells (**Fig. 5a**). Barcode recovery and deep sequencing from each sorted population enabled quantitative, formulation-resolved profiling of cellular distribution across the entire panel in a single experiment. (**Fig. 5b** and **Supplementary Fig. 12**). Heatmap analysis of normalized barcode counts revealed pronounced and formulation-dependent heterogeneity in cellular distribution. Among the screened formulations, HLNP 69 exhibited elevated enrichment in neurons compared with other cell types, HLNP 37 showed preferential enrichment in microglia, HLNP 56 preferentially accumulated in oligodendrocytes, and HLNP 73 exhibited enhanced distribution in endothelial cells. Additional formulations showed comparatively higher enrichment in neurons and astrocytes, highlighting the diverse cellular targeting patterns enabled by the HLNP library.

**Fig. 5:**
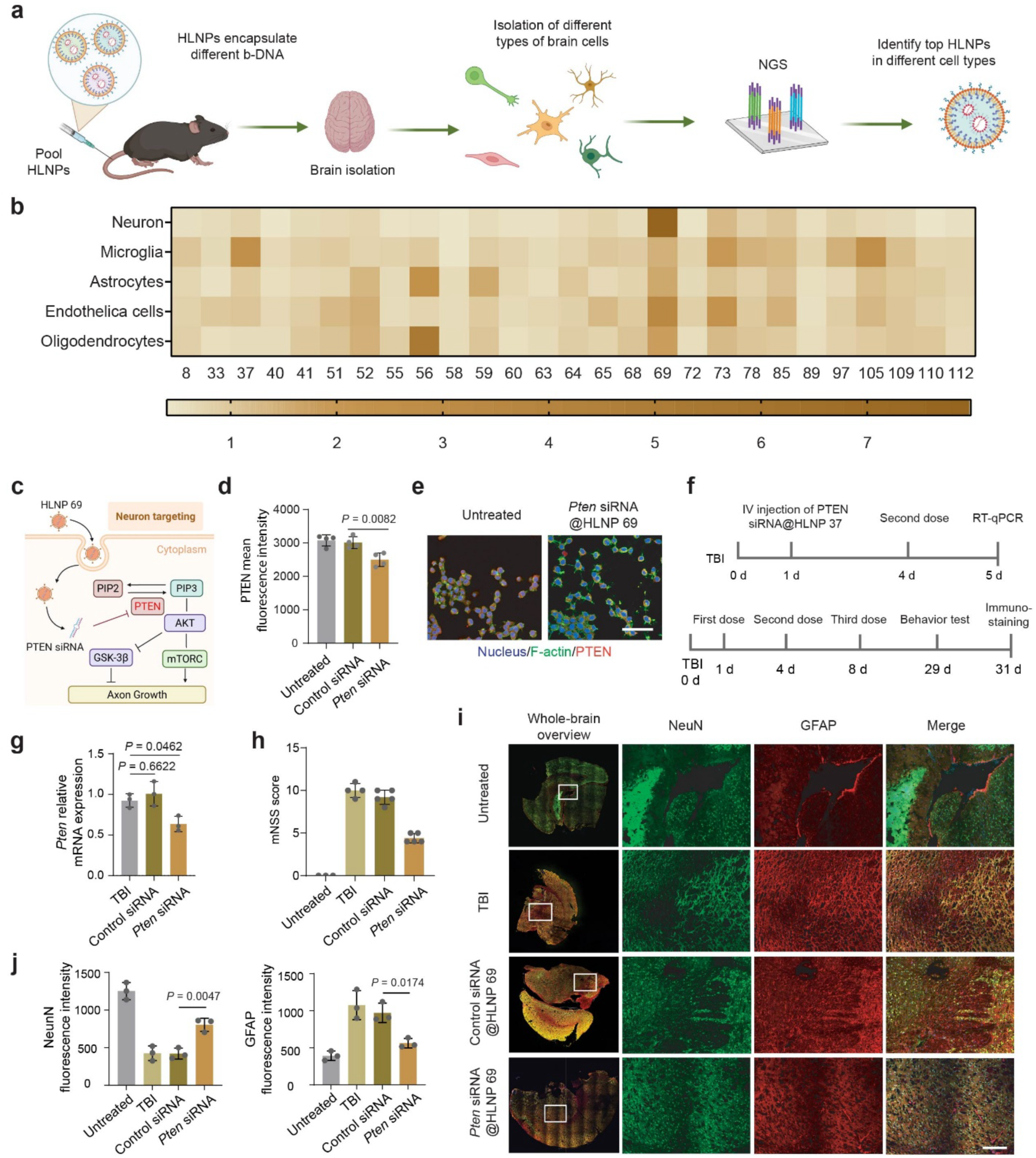
*In vivo* cell-type-resolved screening identifies brain-targeting HLNPs for RNA delivery-mediated neuronal repair and attenuation of neuroinflammation after traumatic brain injury. **a**, Schematic illustration of the *in vivo* DNA-barcoded HLNP screening workflow. Mice were intravenously administered pooled HLNP libraries containing uniquely barcoded formulations. Following systemic circulation, brains were harvested and dissociated into single-cell suspensions, and major brain cell populations were isolated. HLNP-associated DNA barcodes were recovered from each cell population and quantified by NGS to identify formulations enriched in specific brain cell types. **b**, Heatmap showing normalized barcode enrichment of individual HLNP formulations across major brain cell populations, including neurons, microglia, astrocytes, endothelial cells and oligodendrocytes. Color intensity represents the relative enrichment of each formulation within the indicated cell type. Values were independently normalized within each cell type. **c**, Schematic illustration of neuron-targeted delivery of *Pten* siRNA by HLNPs and subsequent activation of the PI3K–AKT signaling pathway to promote axonal growth. **d**, Flow cytometric analysis of PTEN protein expression in N2a cells following treatment with HLNP 69 encapsulating *Pten* siRNA. Data are presented as mean fluorescence intensity (MFI) (*n* = 4). **e**, Representative confocal images of N2a cells stained for nuclei (blue), F-actin (green), and PTEN (red) following treatment with HLNP 69–encapsulated *Pten* siRNA. Scale bar, 50 μm. **f**, Experimental timeline for TBI induction, intravenous administration of HLNP–*Pten* siRNA formulations, behavioral assessment, and downstream molecular and histological analyses. **g**, Relative *Pten* mRNA expression in brain tissue measured by RT–qPCR following treatment with HLNP 69–encapsulated *Pten* siRNA (*n* = 3). **h**, Neurological function assessed using the mNSS at 29 days post-injury (*n* = 3 for Untreated, *n* = 4 for TBI, *n* = 5 for Control siRNA and *Pten* siRNA). **i**, Representative immunofluorescence images of brain sections stained for NeuN and GFAP. Scale bar, 300 μm. **j**, Quantification of NeuN and GFAP fluorescence intensity in brain sections (*n* = 3). Data are presented as mean ± s.d. Statistical significance was determined using one-way ANOVA with appropriate post hoc multiple-comparison tests.

Consistent with these findings, *in vivo* bioluminescence imaging showed stronger brain-associated luminescence signals for formulations 37 and 69 compared with the control formulation Pati (**Supplementary Fig. 13**). Ex vivo imaging of major organs further confirmed enhanced brain expression for HLNPs 37 and 69, whereas the Pati control predominantly produced signals in the liver, consistent with its known liver tropism. Cy5-labeled ssDNA delivery was further evaluated in brain organoids. Three-dimensional fluorescence reconstruction revealed the spatial distribution of Cy5 signals throughout the organoid tissue following treatment with the different LNP formulations (**Supplementary Fig. 14**), indicating efficient nanoparticle penetration into the organoid interior.

We further selected HLNPs 37, 56, 69, 73, and 105 for protein corona analysis. Consistent with observations from the earlier HLNP formulations, these nanoparticles exhibited markedly higher ApoA-I/ApoE and vitronectin/ApoE ratios (**Supplementary Figs. 15,16**). These results further support the potential role of ApoA-I and vitronectin enrichment in facilitating BBB interaction and promoting brain delivery of HLNPs. To functionally validate cell-type selectivity, HLNPs 69 and 37 were selected for further analysis by flow cytometry.

### Cell-type-specific HLNP-mediated RNA delivery promotes neuronal recovery after TBI and suppresses neuroinflammation

To evaluate the therapeutic potential of HLNP-mediated gene silencing *in vivo*, we investigated the delivery of *Pten* siRNA using lead formulation 69, which exhibited the highest neuronal accumulation. PTEN is a key negative regulator of the PI3KAKT signaling pathway whose activity suppresses neuronal survival and axonal regeneration following central nervous system injury, thereby representing a therapeutically relevant target for post-injury intervention (**Fig. 5c, Supplementary Fig. 17**). Prior to *in vivo* evaluation, in vitro validation in N2a neuronal cells confirmed efficient *Pten* knockdown following treatment with *Pten* siRNA@HLNP 69, as quantified by RT-qPCR (**Supplementary Figs. 18**). Consistent with transcript-level suppression, flow cytometry and immunofluorescence imaging revealed a marked reduction in PTEN protein expression in treated cells (**Fig. 5d,e**).

For *in vivo* evaluation, a controlled weight-drop method was employed to induce TBI in mice^41,42^. Following injury, *Pten* siRNA@HLNP 69 was administered intravenously at days 1, 4, and 8 post injury (**Fig. 5f**). At day 5 post injury, RT-qPCR analysis of injured brain tissue revealed a significant reduction in *Pten* mRNA levels in *Pten* siRNA@HLNP treated animals compared with PBS- and control siRNA@HLNP-treated groups (**Fig. 5g**). These molecular alterations were accompanied by improved neurological recovery, as assessed by the modified neurological severity score (mNSS)^43^, a composite behavioral metric evaluating motor, sensory, reflex, and balance functions. Specifically, mice treated with HLNP-*Pten* siRNA exhibited a significant reduction in mNSS at 29 days post-injury (**Fig. 5h**). Immunofluorescence staining further confirmed that *Pten* siRNA treatment preserved neuronal integrity, as evidenced by NeuN staining, and attenuated reactive astrogliosis, evidenced by reduced GFAP signal, compared with untreated or control siRNA@HLNPs–treated TBI mice (**Fig. 5i,j**). Together, these results demonstrate that neuron-targeting HLNP 69 achieves therapeutically meaningful *Pten* silencing *in vivo*, translating molecular target engagement into coordinated cellular and functional improvements after traumatic brain injury.

### Microglia-targeted HLNP 37 delivering IL-10 mRNA suppresses neuroinflammation

Dysregulation of microglial activation is a central driver of neuroinflammatory pathology across a broad spectrum of CNS diseases^44^. To evaluate the therapeutic potential of microglia-targeted RNA delivery, we selected HLNP 37, which exhibited highest microglial enrichment in the cell-type-specific screening, as the delivery vehicle for IL-10 mRNA^45^ in an LPS-induced neuroinflammation model (**Fig. 6a** and **Supplementary Fig. 19**). In vitro validation in LPS-stimulated BV2 microglial cells confirmed that HLNP 37-mediated IL-10 mRNA delivery robustly increased Il10 transcript levels and suppressed proinflammatory gene expression including TNFa and Il1b, with concurrent Socs3 upregulation confirming activation of the IL-10 signaling cascade (**Fig. 6b**). ELISA measurements further confirmed increased IL-10 protein secretion and reduced TNF-alpha production under ongoing LPS stimulation (**Fig. 6c**), demonstrating effective suppression of the inflammatory phenotype at the protein level in vitro.

**Fig. 6:**
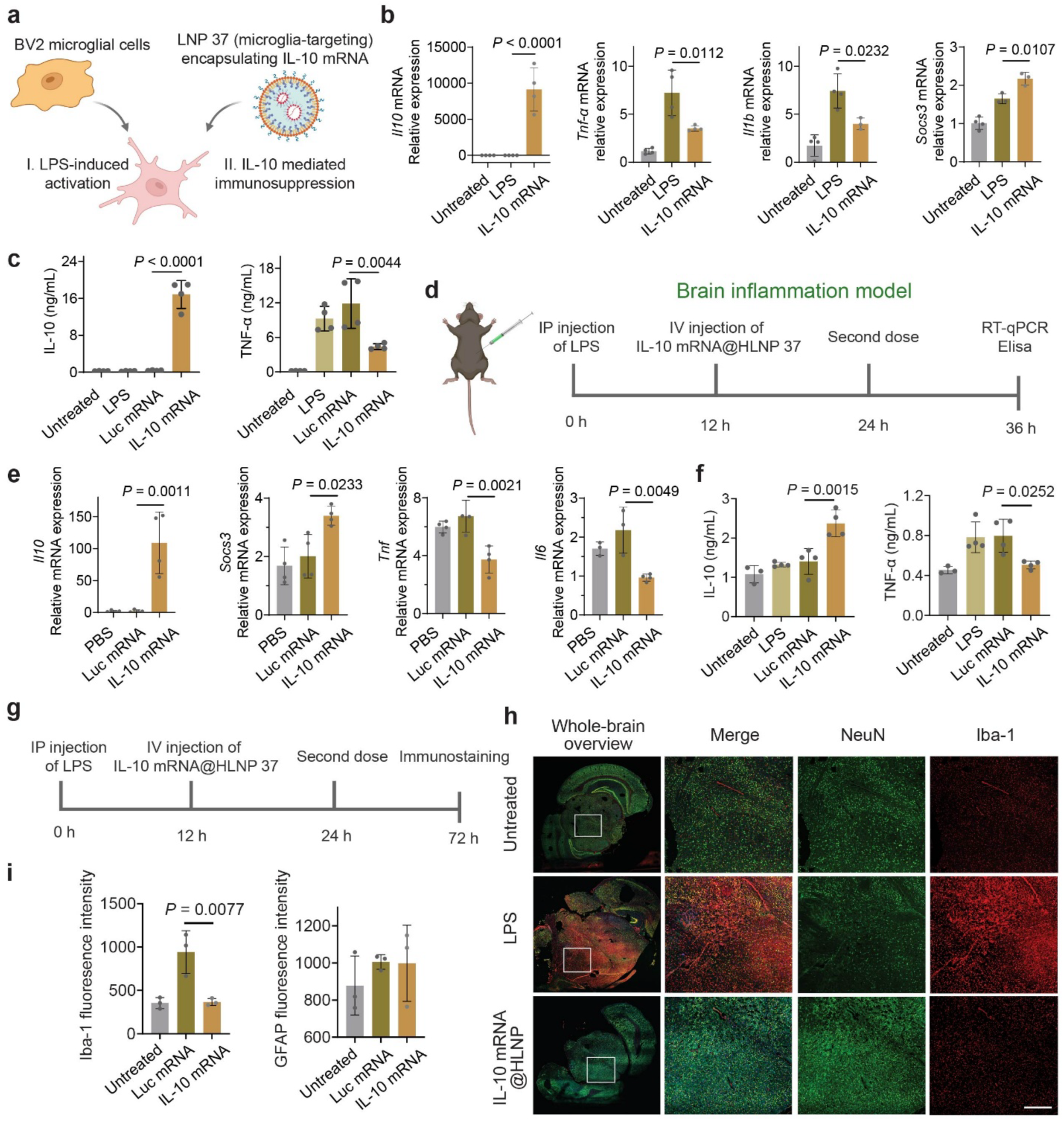
Microglia-targeted HLNP 37-mediated IL-10 mRNA delivery attenuates LPS-induced neuroinflammation. **a**, Schematic illustration of HLNP 37-mediated delivery of IL-10 mRNA to BV2 microglial cells following LPS stimulation. **b**, RT–qPCR analysis of inflammatory gene expression in BV2 cells treated with IL-10 mRNA–loaded HLNP37 following LPS stimulation, including *Il10*, *Tnf*, *Il1b*, and *Socs3* (*n* = 4). **c**, Enzyme-linked immunosorbent assay (ELISA) quantification of IL-10 and TNF-α secretion in BV2 cells after treatment with IL-10 mRNA–loaded HLNP 37 (*n* = 4). **d**, Experimental timeline of the *in vivo* LPS-induced neuroinflammation model. Mice received intraperitoneal (IP) LPS administration followed by intravenous dosing of HLNP 37 encapsulating IL-10 mRNA, with downstream molecular and histological analyses performed at indicated time points. **e**, RT–qPCR quantification of inflammatory gene expression in brain tissue following LPS challenge and HLNP37–IL-10 mRNA treatment, including *Il10, Socs3, Tnf*, and *Il6* (*n* = 4). **f**, ELISA quantification of IL-10 and TNF-α levels in brain tissue homogenates (*n* = 3 for Untreated group, *n* = 4 for other groups). **g**, Experimental timeline of the *in vivo* LPS-induced neuroinflammation model. Mice received intraperitoneal (IP) administration of LPS, followed by intravenous injection of HLNP 37 encapsulating IL-10 mRNA, with downstream molecular and histological analyses performed at the indicated time points. **h**, Representative immunofluorescence images of brain sections stained for NeuN (neuron) and Iba1 (microglia), showing reduced neuroinflammatory activation following HLNP37–IL-10 mRNA treatment in the LPS-induced neuroinflammation model. Scar bar: 50 μm. **i**, Quantification of Iba1 and NeuN fluorescence intensity in brain sections (*n* = 3).

For *in vivo* validation, systemic neuroinflammation was induced by intraperitoneal LPS administration, mice received by two intravenous doses of IL-10 mRNA at 12 and 24 hours post-LPS challenge(**Fig. 6d**). RT-qPCR analysis of brain tissue revealed robust upregulation of *Il10* mRNA (**Fig. 6e**), accompanied by significant downregulation of *Tnf* and *Il6* transcripts, together with increased *Socs3* expression. These results confirmed that the HLNP-delivered IL10 mRNA effectively activated the IL-10-mediated anti-inflammatory signaling pathways within the brain. Consistent with these transcriptional changes, ELISA quantification of brain tissue homogenates demonstrated elevated IL-10 protein levels and a concomitant reduction in TNF-α, confirming functional suppression of neuroinflammatory signaling at the protein level (**Fig. 6f**). Notably, attenuation of TNF-α was observed despite ongoing inflammatory challenge, highlighting the potency and durability of HLNP-mediated IL-10 mRNA expression.

Immunofluorescence staining of brain sections revealed markedly reduced neuroinflammatory signatures in IL-10 mRNA-treated mice compared with LPS-challenged controls (**Fig. 6g-i**). In particular, staining of the microglial marker Iba-1 showed substantially decreased immunoreactivity and reduced accumulation of activated microglia within inflamed regions, consistent with attenuation of microglial activation. Quantification revealed a significant reduction in Iba1 fluorescence intensity following HLNP37-mediated IL-10 mRNA delivery, indicating attenuated microglial activation. In contrast, fluorescence intensity of the neuronal marker NeuN remained largely unchanged across treatment groups. Collectively, these results demonstrate that microglia-targeted HLNP 37 delivery of IL-10 mRNA enables robust, multi-level suppression of LPS-induced neuroinflammation, effectively reprogramming the inflammatory microenvironment toward an anti-inflammatory state both in vitro and *in vivo*.

### Outlook

This work introduces protein corona programming as a design principle for CNS RNA nanomedicine, demonstrating that incorporating HDL-derived phospholipids into LNP formulations biases plasma protein adsorption toward an ApoA-I and vitronectin-enriched corona with reduced ApoE content, redirecting nanoparticle tropism from liver to brain. High-throughput *in vivo* screening of a 112-member HLNP library, combined with quantitative proteomic profiling, established that ApoA-I/ApoE and vitronectin/ApoE corona ratios serve as predictive biomarkers of BBB transport efficiency, linking lipid composition directly to organ-level biodistribution. Cell-type-resolved screening further revealed that lipid composition programs post-BBB cellular distribution in a formulation-dependent manner, enabling selective enrichment across different neural cell types from a single nanoparticle scaffold. Therapeutic validation confirmed that this selectivity translates into meaningful outcomes: neuron-targeted *Pten* siRNA delivery produced functional neurological recovery after TBI, and microglia-targeted IL-10 mRNA delivery suppressed neuroinflammation at transcriptional, translational, and histological levels.

Despite these advances, several important questions remain. First, although our data suggest that ApoA-I enrichment and vitronectin recruitment enhance nanoparticle association with the neurovascular interface, the precise intracellular trafficking routes that enable transcytosis rather than lysosomal degradation require further investigation. Understanding how ApoA-I and vitronectin engage these receptors cooperatively, and whether additional minor corona constituents contribute to BBB crossing, will be important for rational refinement of the HLNP design. Second, the current library explores seven HDL-mimetic phospholipids combined with four cholesterol analogues and four ionizable lipids, but the full formulation space governing corona composition and cell-type targeting remains largely uncharted. Machine learning-guided screening of larger combinatorial libraries, informed by proteomic corona signatures as quantitative design inputs, could accelerate the identification of formulations with higher cell-type specificity and deeper parenchymal penetration. Third, translation to larger animal models and ultimately to humans will require establishing whether the receptor expression patterns that mediate HLNP BBB crossing are conserved across species.

Looking forward, the HLNP platform offers a modular and clinically translatable framework for targeted RNA delivery to the CNS. The principle that engaging endogenous lipoprotein-associated transport pathways represents a promising strategy for overcoming the challenge of brain delivery. More broadly, the HLNP platform opens up possibilities for potential applications in neurodegenerative disease treatment, neuroinflammation modulation, and targeted gene delivery.

## Methods

### Materials

All chemicals were purchased from Fisher Scientific unless otherwise listed. The b-DNA and primers were purchased from Integrated DNA Technologies (IDT). Detailed sequence information is provided in the **Supplementary Information**. The 16:0-18:2 PC (850458), 18:0-18:2 PC (850468), 16:0-20:4 PC (850459), 16:0-18:1 PC (850457), 16:0 SM (D18:1/16:0, 860584), 18:0 SM (D18:1/18:0, 860586), 24:0 SM (860592), and DMG-PEG 2000 (880151) were purchased from Avanti Polar Lipids. SM-102 (33474), DLin-MC3-DMA (34364), ALC-0315 (34337), and C12-200 (36699) were purchased from Cayman Chemical Company. Cholesterol (C8667) and cholesteryl acetate (151114) were purchased from Sigma. Cholesteryl oleate (BP-29625) and cholesterol pelargonate (BP-41508) were purchased from BroadPharm. Animal-Free Recombinant Human Vitronectin was purchased from PeproTech. Native Human Apolipoprotein A1 (0650-0311) was purchased from Bio-Rad. Vitronectin (AF-140-09) was purchased from PeproTech. Origin mouse C57BL6 plasma was purchased from Innovative Research. The FLuc mRNAs was generated using Takara IVTpro T7 mRNA Synthesis Kit. CleanCap EGFP mRNA, mCherry mRNA and IL-10-mRNA were purchased from TriLink. Luciferin (P1043) was purchased from Promega. Pierce Firefly Luciferase Glow Assay Kit (16177) was purchased from Thermo Scientific. BLT-1 (HY-116767), Cilengitide (HY-16141), and Probucol (HY-B0388) were purchased from MedChemExpress. *Ppib* mouse pre-designed siRNA Set was purchased from MedChem Express. *Pten* siRNA was purchased from Horizon Discovery.

### Preparation and characterization of HLNPs

HLNPs encapsulating b-DNA, siRNA, or mRNA were formulated using microfluidic mixing. An ethanol phase containing a defined molar mixture of lipids was rapidly mixed with an aqueous phase (50 mM citrate buffer, pH 4.0) containing the nucleic acid cargo at a flow-rate ratio of 1:3 using automated nanoparticle system Sunshine (Unchained Labs). The HLNPs were prepared at an N/P ratio of 6, defined as the molar ratio of amine groups in the ionizable lipid to phosphate groups in the b-DNA, siRNA, or mRNA. The total lipid concentration ranged from 2 mM to 8 mM. HLNPs were composed of an ionizable lipid, cholesterol, phospholipid, and DMG-PEG2000 at a molar ratio of 50:38.5:10:1.5. HLNPs were dialyzed in 1× PBS using Pur-A-Lyzer Midi Dialysis Kit (6-8 kDa MWCO, Sigma-aldrich) for 3 h. The hydrodynamic diameter and PDI of the resulting HLNPs were measured using a Zetasizer Nano ZS (Malvern Instruments). Encapsulation efficiency was determined using the Quant-iT RiboGreen RNA reagent (R11490, Invitrogen).

### Preparation of the b-DNA library

Each b-DNA consists of single-stranded DNA containing 61 nucleotides with 5 consecutive phosphorothioate bonds at each end. The barcode region (reading region) was composed of 10 nucleotides. An additional 10 random nucleotides were added at the 3’ end of the barcode region to monitor PCR over-amplification. The 5′ and 3′ ends of each b-DNA were conserved and contained priming sites for Illumina adapters. A full list of b-DNA sequences can be found in **Supplementary Table 2**. All oligonucleotides in this study were by Integrated DNA Technologies, unless otherwise specified.

### Animal studies

All animal protocols were approved by the Institutional Animal Care & Use Committee (IACUC) of University of Massachusetts Amherst (Protocol No. 5234), and were consistent with local, state and federal regulations as applicable. C57BL/6J (female, 6-8 weeks, 20-25 g) mice were purchased from Jackson Laboratory. All mice were housed in a specific-pathogen-free animal facility at ambient temperature (22 ± 2 °C), air humidity 40–70% and 12 h dark/12 h light cycle and had free access to water and chow.

### *In vivo* b-DNA–Barcoded HLNP Delivery

To evaluate HLNP biodistribution across multiple tissues, each HLNP formulation carries a distinct b-DNAs. HLNP formulations were organized into multiple pooled groups for each injection and administered intravenously to 8-week-old female C57BL/6 mice (20–25 g).

Intrathecal administration of HLNP formulations was performed in mice to evaluate nanoparticle distribution following direct delivery into the cerebrospinal fluid. Briefly, mice were anesthetized with isoflurane and placed in the prone position. A 30-gauge needle attached to a Hamilton syringe was inserted into the L5–L6 intervertebral space to access the intrathecal compartment. Correct needle placement was confirmed by a characteristic tail-flick response. HLNPs encapsulating b-DNAs were injected slowly in a total volume of 5 μL over ∼10 s. The needle was kept in place for an additional 5 s to minimize reflux before withdrawal. Mice were monitored until full recovery from anesthesia and returned to their home cages.

### Tissue Dissociation and b-DNA Extraction

Four hours post-injection, tissues (brain, heart, liver, spleen, lung, and kidney) were harvested. Brain tissues were dissociated using the Adult Brain Dissociation Kit (130-107-677, Miltenyi Biotec) according to the manufacturer’s instructions. Briefly, each brain was transferred into a gentleMACS C Tube containing the provided enzyme mix and subjected to combined enzymatic and mechanical dissociation using a gentleMACS Octo Dissociator (Miltenyi Biotec). The homogenate was filtered through a 70 µm cell strainer (130-110-916, Miltenyi Biotec) to obtain a single-cell suspension. Debris was removed using Debris Removal Solution (130-109-398, Miltenyi Biotec). Residual erythrocytes were lysed using Red Blood Cell Lysis Solution (130-094-183, Miltenyi Biotec). Cells were washed and resuspended in 80 μL of DPBS containing 0.5% bovine serum albumin for each brain homogenate.

Peripheral organs (heart, liver, spleen, lung, and kidney) were dissociated using the Multi Tissue Dissociation Kit (130-110-204, Miltenyi Biotec) in combination with a gentleMACS Octo Dissociator.

For b-DNA extraction, samples were incubated in QuickExtract DNA Extraction Solution (Biosearch Technologies) at 65 °C for 6 min, vortexed briefly (15 s), followed by incubation at 98 °C for 2 min. Lysates were cooled to room temperature (∼5 min) and stored at −20 °C until further use.

### Brain Cell-Type–Specific Analysis

For cell-type–specific analyses, single-cell suspensions were processed immediately after tissue dissociation. For each sample, cells from two mice were pooled prior to magnetic-activated cell sorting (MACS). Neural cell populations were enriched by MACS (Miltenyi Biotec) using the following kits according to the manufacturer’s instructions: Neuron Isolation Kit, mouse (130-115-389); CD11b (microglia) MicroBeads (130-093-634); Anti-ACSA-2 MicroBead Kit (130-097-678); Anti-O4 MicroBeads (130-094-543); and CD31 MicroBeads (130-097-418). Following isolation, cells were lysed and b-DNA was extracted using the QuickExtract protocol described above.

### b-DNA amplification and sequencing library preparation

Following DNA extraction from tissues, samples were subjected to PCR amplification. Reactions (25 μL total volume) were prepared using 12.5 μL KAPA HotStart ReadyMix (KK2602, Roche), 2.5 μL of 10× UDG buffer, 1 μL UDG enzyme, 2.5 μL each of forward and reverse primers (5 μM), 2.5 μL template DNA, and nuclease-free water to volume.

PCR cycling conditions consisted of an initial denaturation at 98 °C for 3 min, followed by 28 cycles of 95 °C for 15 s, 62 °C for 15 s, and 72 °C for 20 s, with a final extension at 72 °C for 1 min and a hold at 4 °C. PCR products were analyzed by 4% agarose gel electrophoresis and stored at −20 °C until further processing.

PCR products were pooled and purified using AMPure XP beads (Beckman Coulter). Briefly, DNA was bound to beads, washed twice with 70% ethanol, and eluted in low-salt TE buffer. Purified DNA was stored at −20 °C until sequencing.

### Deep sequencing and delivery quantification

Deep sequencing was performed using Illumina next-generation sequencing, with libraries sequenced on an Illumina MiSeq platform. Sequencing data were analyzed using a custom Python algorithm to quantify reads corresponding to each b-DNA across tissues.

For both intravenous and intrathecal injections, samples were organized into three groups, each containing an internal HLNP control. Within each group, read counts were first normalized to the corresponding internal HLNP based on sequencing reads. The relative delivery of each HLNP formulation to a given organ was calculated by dividing the corresponding b-DNA read count by the total b-DNA reads recovered from that tissue, yielding a normalized biodistribution value.

For cell-type–specific analysis, the number of reads of a given b-DNA in a specific cell type was divided by the total number of b-DNA reads detected in the brain. This normalization allowed assessment of cell-type–level biodistribution and facilitated identification of the lead HLNP formulations for each target cell type.

### In vitro luciferase assay for HLNPs

N2a, BV2 and bEnd.3 cells were cultured in Dulbecco’s modified Eagle’s medium (DMEM) supplemented with 10% fetal bovine serum and 1% penicillin–streptomycin at 37 °C in a humidified atmosphere with 5% CO₂. Cells were seeded into white 96-well plates at a density of 1 × 10⁴ cells per well and allowed to adhere overnight. HLNPs encapsulating FLuc mRNA were added to each well (20 ng per well). After 24 h incubation, luciferase expression was measured using Bright-Glo luciferase substrate. Luminescence was recorded 5 min after substrate addition using a BioTek multimode plate reader.

### In vitro mcherry reporter assay for HLNPs

N2a and BV2 cells were separately seeded into 96-well plates at a density of 1 × 10⁴ cells per well and allowed to adhere overnight. HLNPs encapsulating mRNA encoding mCherry were added to each well (200 ng per well). After 24 h incubation, mCherry expression was imaged using a CellCyte X live-cell imaging system (Cytena).

### RT-qPCR analysis

N2a cells were seeded in 24-well plates at a density of 5 × 10⁴ cells per well and cultured overnight in DMEM supplemented with 10% FBS and 1% penicillin–streptomycin. Cells were treated with HLNP formulations encapsulating *Ppib* siRNA or *Pten* siRNA at a final concentration of 50 nM and incubated for 24 h prior to downstream analysis.

BV2 microglial cells were seeded in 24-well plates at a density of 5 × 10⁴ cells per well and cultured overnight in DMEM supplemented with 10% FBS and 1% penicillin–streptomycin. Cells were stimulated with LPS (100 ng mL⁻¹) prior to or during nanoparticle treatment as indicated, followed by incubation with HLNP formulations encapsulating IL-10 mRNA (20 ng per well) for 4 h.

Brain tissues were collected at designated time points, rapidly dissected, and transferred into TRIzol reagent. Samples were homogenized on ice using a bead-based tissue homogenizer with stainless steel beads until complete tissue disruption was achieved.

Total RNA was extracted using TRIzol reagent according to the manufacturer’s protocol. RNA was further purified using the Monarch Total RNA Miniprep Kit (T2010S, New England Biolabs). RNA concentration and purity were measured using a NanoDrop spectrophotometer. RT–qPCR was performed using the Luna Universal One-Step RT–qPCR Kit (New England Biolabs) on a CFX96 Real-Time PCR system (Bio-Rad). Relative gene expression levels were normalized to the housekeeping gene *Gapdh* and calculated using the 2^−ΔΔCT^ method.

### Protein Corona Analysis

IOLNPs were prepared as described above and incubated with pooled mouse serum to allow protein corona formation^38^. Briefly, mouse serum was incubated with IOLNPs at 37 °C for 1 h under gentle agitation. After incubation, nanoparticle–protein complexes were isolated by magnetic separation using a MACS Cell Separation system. IOLNPs were retained on MACS columns, while unbound proteins were removed by extensive washing; the bound protein corona fraction was subsequently eluted. Protein concentration was quantified using a BCA assay. Protein corona samples were subjected to in-solution digestion using standard reduction, alkylation, and tryptic digestion procedures, followed by C18 Stage-Tip cleanup. The trypsin digest was analyzed by nanoLC-MS/MS at the University of Massachusetts Amherst Mass Spectrometry Core Facility (RRID:SCR_019063). Analyses were performed using an Orbitrap Fusion mass spectrometer coupled to an Easy-nLC 1000 nanoLC chromatography system. Mass spectral data were processed using Proteome Discoverer version 2.5 software (Thermo Scientific).

### In vitro BBB transwell assay

An in vitro BBB model was established using a Transwell co-culture system. Briefly, bEnd.3 mouse brain endothelial cells were seeded onto the upper chamber of Transwell inserts (0.4 µm pore size) pre-coated with collagen and allowed to form a confluent monolayer over 10 days. N2a cells were seeded in the lower chamber and cultured to ∼70% confluence. HLNPs were pre-incubated with ApoA-I or vitronectin, at a mass ratio of 10:1 (Lipid : protein) for 1 h at 37 °C to allow protein corona formation. HLNPs encapsulating Fluc mRNA (200 ng per well) were then added to the apical (upper) chamber and incubated for 24 h. Following incubation, luciferase expression in N2a and bEnd.3 cells were quantified using the Bright-Glo luciferase assay system according to the manufacturer’s instructions.

### In vitro receptor inhibition assay

To investigate the involvement of specific receptors in HLNP uptake, receptor inhibition studies were performed using pharmacological inhibitors. bEnd.3 cells were seeded in white 96-well plates at a density of 1 × 10⁴ cells per well. HLNPs were pre-incubated with 10% (v/v) mouse plasma at 37 °C for 1 h to promote the formation of a protein corona. Prior to nanoparticle treatment, cells were pre-treated for 1 h at 37 °C with either the SR-BI inhibitor BLT-1 (10 μM) or the integrin αvβ3/αvβ5 inhibitor cilengitide (5 μM). HLNPs were then added to cells in the continued presence of inhibitors without removal and incubated for 24 h, after which luciferase expression was quantified.

### HLNP-FLuc mRNA delivery by intravenous administration via tail vein

C57BL/6J mice were intravenously injected with selected HLNPs (*N*^1^-methyl-pseudouridine FLuc mRNA, 0.5 mg kg^−1^). After 6 h, the mice received an intraperitoneal injection of 150 μl of d-luciferin substrate (30 mg ml^−1^). The major organs, including the brain, were dissected and imaged after 8 min using a Xenogen IVIS imaging system. The signals in the brain of all the groups were quantified using regions of interest.

### Flow cytometry analysis of PTEN knockdown

N2a cells were seeded in 24-well plates at a density of 5 × 10⁴ cells per well and cultured overnight. Cells were treated with HLNP formulations encapsulating PTEN siRNA (50 nM) for 36 h. Cells were then harvested by trypsinization, washed twice with PBS, and fixed with 4% paraformaldehyde for 10–15 min at room temperature. After permeabilization with 0.1% Triton X-100 and blocking with 1% BSA, cells were incubated with anti-PTEN primary antibody (1:200) for 1 h at room temperature, followed by incubation with fluorophore-conjugated secondary antibody (1:500) for 30 min in the dark. Cells were washed and resuspended in PBS for analysis on a flow cytometer (BD LSRFortessa). Mean fluorescence intensity (Mean FI) of PTEN staining was quantified using FlowJo and normalized to untreated or control siRNA–treated cells.

### Immunofluorescence staining and confocal imaging of PTEN expression

N2a cells were seeded on glass coverslips in 24-well plates at a density of 5 × 10⁴ cells per well and cultured overnight. Following treatment with HLNPs encapsulating *Pten* siRNA (50 nM) for 36 h, cells were fixed with 4% paraformaldehyde for 15 min, washed with PBS, and permeabilized with 0.1% Triton X-100 for 10 min. After blocking with 3% BSA for 1 h, cells were incubated with anti-PTEN primary antibody (1:200, PTEN (D4.3) rabbit monoclonal antibody, #9188, Cell Signaling Technology) overnight at 4 °C, followed by incubation with fluorophore-conjugated secondary antibody (1:500, Donkey Anti-Rabbit IgG H&L (Alexa Fluor 568), ab175470, Abcam) for 1 h at room temperature in the dark. Nuclei were counterstained with Hoechst 33342 (1 μg mL⁻¹) for 10 min. Coverslips were mounted and imaged using a Nikon AXR-NSPARC scanning confocal microscope.

### Weight-drop TBI models

A weight-drop TBI model with moderate or severe injury was established according to previously reported procedures^42^. Briefly, brain contusion was induced by free-fall impact. Healthy C57BL/6 mice (8 weeks old, ∼20 g) were anesthetized with isoflurane, a 50 g weight was positioned 70 cm above the head and released along a vertical guide tube, allowing it to freely fall and impact the right hemisphere through a 3 mm diameter impact tip. After injury, mice were placed on a 30 °C heating pad until fully recovered from anesthesia. *Pten* siRNA-loaded HLNPs were administered via intravenous injection through the tail vein. The first dose was given at 1 day post-injury, followed by additional doses on days 4 and 8. For each administration, LNPs were freshly prepared and diluted in sterile PBS prior to injection.

### Behavior analysis

The mNSS was performed as previously described^43^. Prior to testing, mice were habituated to handling by tail suspension (30 s per day for 3 consecutive days) and acclimated to the behavioral apparatus (10 min on beam and open field) 24 h before baseline assessment. Neurological function was evaluated using the modified neurological severity score (mNSS), which integrates motor, sensory, balance, and reflex tests. The mNSS battery included: (i) tail suspension (30 s) to assess forelimb and hindlimb flexion and head deviation (≥10°); (ii) open-field walking in a flat arena to evaluate gait abnormalities such as circling or falling; (iii) sensory placing via a visual–tactile forepaw placement test; (iv) proprioceptive response assessed by paw withdrawal at the edge of a table; (v) beam balance on a 100 cm × 2 cm beam for 60 s, scored on a scale of 0–6; and (vi) reflex testing, including pinna, corneal, and startle (paper snap) reflexes, along with observation for dystonia or abnormal movements. Each task was scored according to established mNSS criteria, with higher scores indicating greater neurological impairment. All sub-scores were independently recorded by two investigators blinded to the experimental groups using a stopwatch and standardized data sheets, yielding a composite score ranging from 0 (normal) to 18 (maximal deficit).

### LPS-Induced Brain Inflammation Model and HLNP Treatment

An acute neuroinflammation model was established by systemic administration of lipopolysaccharide (LPS, O111:B4). Eight-week-old female C57BL/6 mice were randomly assigned to experimental groups. At 0 h, mice received an intraperitoneal (i.p.) injection of LPS (1 mg kg⁻¹ in sterile PBS) to induce systemic inflammation and subsequent neuroinflammation, while control animals received an equal volume of PBS. At 12 h post-LPS injection, mice were intravenously administered HLNP 37 encapsulating IL-10 mRNA via tail vein injection (1 mg kg⁻¹ mRNA in 200 µL sterile PBS). A second intravenous dose of the same formulation was administered at 24 h.

### ELISA

The concentrations of tumor necrosis factor-α (TNF-α) and interleukin-6 (IL-6) in serum or tissue homogenates were quantified using commercially available ELISA kits according to the manufacturer’s instructions. For tissue samples, freshly collected tissues were weighed and homogenized in ice-cold RIPA buffer containing protease inhibitors (1:10, w/v), followed by centrifugation at 12,000 × g for 10 min at 4 °C. The supernatants were collected, and total protein concentrations were determined using a BCA assay. Standards and samples were added in duplicate to pre-coated 96-well plates and incubated with detection antibodies and HRP-conjugated streptavidin according to standard procedures. TMB substrate was used for color development, and absorbance was measured at 450 nm.

### Immunofluorescence Staining of Brain Sections

At the indicated time point, mice were deeply anesthetized and transcardially perfused with PBS, followed by 4% paraformaldehyde (PFA). Brains were harvested, post-fixed overnight in 4% PFA at 4 °C, cryoprotected in 30% sucrose, embedded in OCT compound, and sectioned at 10 µm thickness using a cryostat.

For immunofluorescence staining, brain sections were permeabilized with 0.3% Triton X-100 in PBS for 15 min and blocked with 5% normal goat serum in PBS for 1 h at room temperature to minimize nonspecific binding. Sections were then incubated overnight at 4 °C with primary antibodies diluted in blocking buffer. For the TBI model, sections were stained with anti-NeuN (ab104224, Abcam) and anti-GFAP (ab7260, Abcam) antibodies to assess neuronal integrity and astrocyte activation. For the neuroinflammation model, sections were stained with anti-NeuN (ab104224, Abcam) and anti-Iba1 antibody (ab178846, Abcam) antibodies to evaluate neuronal apoptosis.

Following primary antibody incubation, sections were washed three times with PBS (5 min each) and incubated with the appropriate fluorophore-conjugated secondary antibodies diluted in blocking buffer for 1 h at room temperature in the dark. After secondary antibody incubation, sections were washed three times with PBS (5 min each), and nuclei were counterstained with DAPI for 5 min. Fluorescence images were acquired using a fluorescence microscope or confocal microscope. Quantification of fluorescence intensity or the number of positive cells was performed using ImageJ.

### Statistics and reproducibility

Statistical analysis was performed using GraphPad Prism, ImageJ and OriginPro software. Data are presented as mean ± s.d. of biological replicates unless otherwise stated. Detailed statistical methods are described in the main text and figure legends.

## Acknowledgements

This study was supported by Alzheimer’s Association Research Grants (AARG-25-1487957) and AHA Career Award (26CDA1611739). We thank Dr. Chuang Liu at National University of Singapore for technical assistance with IVT-based mRNA synthesis. We acknowledge the UMass Amherst Institute for Applied Life Sciences (IALS) Mass Spectrometry Center and Dr. Steve Eyles for assistance with protein corona characterization and analysis. We thank Dr. Amy Burnside at the Flow Cytometry Core Facility for training and helpful discussions. We also thank Dr. James Chambers and the staff of the IALS Light Microscopy Facility for their guidance and technical support. We thank Jennifer Ser-Dolansky and Sallie Schneider at UMass Amherst for providing histological sectioning services. We acknowledge the BPF Genomics Core Lab at Harvard Medical School for sequencing services. Some figures were created with BioRender.com.

## Contributions

J.G. and Z.X conceived and designed the study. Z.X. performed the experiments and analyzed the data. K.S. assisted with nanoparticle formulation, characterization and animal experiments. V.D. and A.Y. assisted with barcode recovery and sequencing data analysis. Z.X. wrote the manuscript with input from all authors. J.G. supervised the project.

## Ethics declarations/Competing interests

The authors declare no competing interests.

## References

1 Feigin, V. L., Nichols, E., Alam, T., Bannick, M. S., Beghi, E., Blake, N., Culpepper, W. J., Dorsey, E. R., Elbaz, A., Ellenbogen, R. G., Fisher, J. L., Fitzmaurice, C., Giussani, G., Glennie, L., James, S. L. et al. Global, regional, and national burden of neurological disorders, 1990-2016: a systematic analysis for the Global Burden of Disease Study 2016. The Lancet Neurology 18, 459–480 (2019).

2 Wang, S., Weissman, D. & Dong, Y. RNA chemistry and therapeutics. Nature Reviews Drug Discovery 24, 828–851 (2025).

3 Fire, A., Xu, S., Montgomery, M. K., Kostas, S. A., Driver, S. E. & Mello, C. C. Potent and specific genetic interference by double-stranded RNA in Caenorhabditis elegans. Nature 391, 806–811 (1998).

4 Sahin, U., Karikó, K. & Türeci, Ö. mRNA-based therapeutics — developing a new class of drugs. Nature Reviews Drug Discovery 13, 759–780 (2014).

5 Damase, T. R., Sukhovershin, R., Boada, C., Taraballi, F., Pettigrew, R. I. & Cooke, J. P. The Limitless Future of RNA Therapeutics. Frontiers in Bioengineering and Biotechnology **Volume** 9 -2021 (2021).

6 Hu, B., Zhong, L., Weng, Y., Peng, L., Huang, Y., Zhao, Y. & Liang, X.-J. Therapeutic siRNA: state of the art. Signal Transduction and Targeted Therapy 5, 101 (2020).

7 Whitehead, K. A., Langer, R. & Anderson, D. G. Knocking down barriers: advances in siRNA delivery. Nature Reviews Drug Discovery 8, 129–138 (2009).

8 Pardridge, W. M. The blood-brain barrier: Bottleneck in brain drug development. NeuroRX 2, 3–14 (2005).

9 Abbott, N. J., Patabendige, A. A. K., Dolman, D. E. M., Yusof, S. R. & Begley, D. J. Structure and function of the blood–brain barrier. Neurobiology of Disease 37, 13–25 (2010).

10 Zeisel, A., Hochgerner, H., Lönnerberg, P., Johnsson, A., Memic, F., van der Zwan, J., Häring, M., Braun, E., Borm, L. E., La Manno, G., Codeluppi, S., Furlan, A., Lee, K., Skene, N., Harris, K. D. et al. Molecular Architecture of the Mouse Nervous System. Cell 174, 999–1014.e1022 (2018).

11 Manzari, M. T., Shamay, Y., Kiguchi, H., Rosen, N., Scaltriti, M. & Heller, D. A. Targeted drug delivery strategies for precision medicines. Nature Reviews Materials 6, 351–370 (2021).

12 Adams, D., Gonzalez-Duarte, A., O’Riordan William, D., Yang, C.-C., Ueda, M., Kristen Arnt, V., Tournev, I., Schmidt Hartmut, H., Coelho, T., Berk John, L., Lin, K.-P., Vita, G., Attarian, S., Planté-Bordeneuve, V., Mezei Michelle, M. et al. Patisiran, an RNAi Therapeutic, for Hereditary Transthyretin Amyloidosis. New England Journal of Medicine 379, 11–21 (2018).

13 Polack Fernando, P., Thomas Stephen, J., Kitchin, N., Absalon, J., Gurtman, A., Lockhart, S., Perez John, L., Pérez Marc, G., Moreira Edson, D., Zerbini, C., Bailey, R., Swanson Kena, A., Roychoudhury, S., Koury, K., Li, P., et al. Safety and Efficacy of the BNT162b2 mRNA Covid-19 Vaccine. New England Journal of Medicine 383, 2603–2615 (2020).

14 Akinc, A., Querbes, W., De, S., Qin, J., Frank-Kamenetsky, M., Jayaprakash, K. N., Jayaraman, M., Rajeev, K. G., Cantley, W. L., Dorkin, J. R., Butler, J. S., Qin, L., Racie, T., Sprague, A., Fava, E. et al. Targeted Delivery of RNAi Therapeutics With Endogenous and Exogenous Ligand-Based Mechanisms. Molecular Therapy 18, 1357–1364 (2010).

15 Sebastiani, F., Yanez Arteta, M., Lerche, M., Porcar, L., Lang, C., Bragg, R. A., Elmore, C. S., Krishnamurthy, V. R., Russell, R. A., Darwish, T., Pichler, H., Waldie, S., Moulin, M., Haertlein, M., Forsyth, V. T. et al. Apolipoprotein E Binding Drives Structural and Compositional Rearrangement of mRNA-Containing Lipid Nanoparticles. ACS Nano 15, 6709–6722 (2021).

16 Wang, C., Xue, Y., Markovic, T., Li, H., Wang, S., Zhong, Y., Du, S., Zhang, Y., Hou, X., Yu, Y., Liu, Z., Tian, M., Kang, D. D., Wang, L., Guo, K. et al. Blood–brain-barrier-crossing lipid nanoparticles for mRNA delivery to the central nervous system. Nature Materials 24, 1653–1663 (2025).

17 Liu, Z., Zhang, Y., Li, H., Guo, K., Tian, M., Cao, D., Kang, D. D., Xue, Y., Hou, X., Wang, C., Wang, S., Zhong, Y., Yu, C., Deng, B., McComb, D. W. et al. Furan-Derived Lipid Nanoparticles for Transporting mRNA to the Central Nervous System. Journal of the American Chemical Society 147, 16007–16017 (2025).

18 Johnsen, K. B., Gudbergsson, J. M., Skov, M. N., Pilgaard, L., Moos, T. & Duroux, M. A comprehensive overview of exosomes as drug delivery vehicles — Endogenous nanocarriers for targeted cancer therapy. Biochimica et Biophysica Acta (BBA) - Reviews on Cancer 1846, 75–87 (2014).

19 Pardridge, W. M. Blood-Brain Barrier and Delivery of Protein and Gene Therapeutics to Brain. Frontiers in Aging Neuroscience **Volume** 11 - 2019 (2020).

20 Corbo, C., Molinaro, R., Taraballi, F., Toledano Furman, N. E., Hartman, K. A., Sherman, M. B., De Rosa, E., Kirui, D. K., Salvatore, F. & Tasciotti, E. Unveiling the in Vivo Protein Corona of Circulating Leukocyte-like Carriers. ACS Nano 11, 3262–3273 (2017).

21 Monopoli, M. P., Åberg, C., Salvati, A. & Dawson, K. A. Biomolecular coronas provide the biological identity of nanosized materials. Nature Nanotechnology 7, 779–786 (2012).

22 Feng, Y., Tai, W., Huang, P., Qi, S., Yu, X., Li, M., Li, M., Zhang, M., Cao, F., Gao, X., Yang, K., Bai, B., Lei, J., Cheng, M., Li, Y. et al. Albumin-recruiting lipid nanoparticle potentiates the safety and efficacy of mRNA vaccines by avoiding liver accumulation. Nature Materials 24, 1826–1839 (2025).

23 Lundqvist, M., Stigler, J., Elia, G., Lynch, I., Cedervall, T. & Dawson, K. A. Nanoparticle size and surface properties determine the protein corona with possible implications for biological impacts. Proceedings of the National Academy of Sciences 105, 14265–14270 (2008).

24 Voke, E., Arral, M. L., Squire, H. J., Lin, T.-J., Zheng, L., Coreas, R., Lui, A., Iavarone, A. T., Pinals, R. L., Whitehead, K. A. & Landry, M. P. Protein corona formed on lipid nanoparticles compromises delivery efficiency of mRNA cargo. Nature Communications 16, 8699 (2025).

25 Rhea, E. M. & Banks, W. A. Interactions of Lipids, Lipoproteins, and Apolipoproteins with the Blood-Brain Barrier. Pharmaceutical Research 38, 1469–1475 (2021).

26 Mahley, R. W. Apolipoprotein E: from cardiovascular disease to neurodegenerative disorders. Journal of Molecular Medicine 94, 739–746 (2016).

27 von Eckardstein, A., Nordestgaard, B. G., Remaley, A. T. & Catapano, A. L. High-density lipoprotein revisited: biological functions and clinical relevance. European Heart Journal 44, 1394–1407 (2023).

28 Kontush, A., Lhomme, M. & Chapman, M. J. Unraveling the complexities of the HDL lipidome1. Journal of Lipid Research 54, 2950–2963 (2013).

29 Liang, S., Gao, S., Fu, S., Yuan, S., Liu, J., Liang, M., Han, L., Zhang, Z., Liu, Y. & Zhang, N. Screening Natural Cholesterol Analogs to Assemble Self-Adjuvant Lipid Nanoparticles for Antigens Tagging Guided Therapeutic Tumor Vaccine. Advanced Materials 37, 2419182 (2025).

30 Li, B., Jiang, A. Y., Raji, I., Atyeo, C., Raimondo, T. M., Gordon, A. G. R., Rhym, L. H., Samad, T., MacIsaac, C., Witten, J., Mughal, H., Chicz, T. M., Xu, Y., McNamara, R. P., Bhatia, S. et al. Enhancing the immunogenicity of lipid-nanoparticle mRNA vaccines by adjuvanting the ionizable lipid and the mRNA. Nature Biomedical Engineering 9, 167–184 (2025).

31 Swingle, K. L., Hamilton, A. G., Safford, H. C., Geisler, H. C., Thatte, A. S., Palanki, R., Murray, A. M., Han, E. L., Mukalel, A. J., Han, X., Joseph, R. A., Ghalsasi, A. A., Alameh, M.-G., Weissman, D. & Mitchell, M. J. Placenta-tropic VEGF mRNA lipid nanoparticles ameliorate murine pre-eclampsia. Nature 637, 412–421 (2025).

32 Patel, S., Ashwanikumar, N., Robinson, E., Xia, Y., Mihai, C., Griffith, J. P., Hou, S., Esposito, A. A., Ketova, T., Welsher, K., Joyal, J. L., Almarsson, Ö. & Sahay, G. Naturally-occurring cholesterol analogues in lipid nanoparticles induce polymorphic shape and enhance intracellular delivery of mRNA. Nature Communications 11, 983 (2020).

33 Eygeris, Y., Patel, S., Jozic, A. & Sahay, G. Deconvoluting Lipid Nanoparticle Structure for Messenger RNA Delivery. Nano Letters 20, 4543–4549 (2020).

34 Paunovska, K., Gil, C. J., Lokugamage, M. P., Sago, C. D., Sato, M., Lando, G. N., Gamboa Castro, M., Bryksin, A. V. & Dahlman, J. E. Analyzing 2000 in Vivo Drug Delivery Data Points Reveals Cholesterol Structure Impacts Nanoparticle Delivery. ACS Nano 12, 8341–8349 (2018).

35 Zenhausern, R., Jang, B., Schrader Echeverri, E., Gentry, K., Calkins, R., Curran, E. H., Wood, J. S., Stammen, R. L., Loughrey, D., Chappa, P., Koveal, D., Kim, H. & Dahlman, J. E. Lipid nanoparticle screening in nonhuman primates with minimal loss of life. Nature Biotechnology (2025).

36 Vanni, S., Riccardi, L., Palermo, G. & De Vivo, M. Structure and Dynamics of the Acyl Chains in the Membrane Trafficking and Enzymatic Processing of Lipids. Accounts of Chemical Research 52, 3087–3096 (2019).

37 Akinc, A., Maier, M. A., Manoharan, M., Fitzgerald, K., Jayaraman, M., Barros, S., Ansell, S., Du, X., Hope, M. J., Madden, T. D., Mui, B. L., Semple, S. C., Tam, Y. K., Ciufolini, M., Witzigmann, D. et al. The Onpattro story and the clinical translation of nanomedicines containing nucleic acid-based drugs. Nature Nanotechnology 14, 1084–1087 (2019).

38 Francia, V., Zhang, Y., Cheng, M. H. Y., Schiffelers, R. M., Witzigmann, D. & Cullis, P. R. A magnetic separation method for isolating and characterizing the biomolecular corona of lipid nanoparticles. Proceedings of the National Academy of Sciences 121, e2307803120 (2024).

39 Hosseini-Kharat, M., Bremmell, K. E. & Prestidge, C. A. Why do lipid nanoparticles target the liver? Understanding of biodistribution and liver-specific tropism. Molecular Therapy Methods & Clinical Development 33, 101436 (2025).

40 Aliyandi, A., Reker-Smit, C., Zuhorn, I. S. & Salvati, A. Cell surface biotinylation to identify the receptors involved in nanoparticle uptake into endothelial cells. Acta Biomaterialia 155, 507–520 (2023).

41 Mannix, R., Meehan, W. P., Mandeville, J., Grant, P. E., Gray, T., Berglass, J., Zhang, J., Bryant, J., Rezaie, S., Chung, J. Y., Peters, N. V., Lee, C., Tien, L. W., Kaplan, D. L., Feany, M. et al. Clinical correlates in an experimental model of repetitive mild brain injury. Annals of Neurology 74, 65–75 (2013).

42 Kondo, A., Shahpasand, K., Mannix, R., Qiu, J., Moncaster, J., Chen, C.-H., Yao, Y., Lin, Y.-M., Driver, J. A., Sun, Y., Wei, S., Luo, M.-L., Albayram, O., Huang, P., Rotenberg, A. et al. Antibody against early driver of neurodegeneration cis P-tau blocks brain injury and tauopathy. Nature 523, 431–436 (2015).

43 Chen, J., Sanberg, P. R., Li, Y., Wang, L., Lu, M., Willing, A. E., Sanchez-Ramos, J. & Chopp, M. Intravenous Administration of Human Umbilical Cord Blood Reduces Behavioral Deficits After Stroke in Rats. Stroke 32, 2682–2688 (2001).

44 Gao, C., Jiang, J., Tan, Y. & Chen, S. Microglia in neurodegenerative diseases: mechanism and potential therapeutic targets. Signal Transduction and Targeted Therapy 8, 359 (2023).

45 Gao, M., Li, Y., Ho, W., Chen, C., Chen, Q., Li, F., Tang, M., Fan, Q., Wan, J., Yu, W., Xu, X., Li, P. & Zhang, X.-Q. Targeted mRNA Nanoparticles Ameliorate Blood–Brain Barrier Disruption Postischemic Stroke by Modulating Microglia Polarization. ACS Nano 18, 3260–3275 (2024).

