## Supporting information for "High-density Lipoprotein Inspired Lipid Nanoparticles for Systemic RNA delivery to the Brain"

**Table of Contents**

Supplementary Materials

Supplementary Methods

Supplementary Figures

Supplementary Tables

**Materials**

Quant-iT™ RiboGreen™ RNA Assay Kit (R11490) was purchased from Thermo Scientifi. Chlorpromazine (hydrochloride) (16129), Genistein (10005167), 5-(N-ethyl-N-isopropyl)-Amiloride (14406) and Cytochalasin D (11330) were purchased from Cayman Chemical Company. Methyltetrazine-amine HCl salt (BP-22433) was purchased from BroadPharm. Dynasore (HY-15304) was purchased from MedChem Express. 5-(N-Ethyl-N-isopropyl)-amiloride (A3085), MβCD (C4555) and chlorpromazine (1125006) were purchased from Sigma-Aldrich. Organoids were kindly provided by Dr. Yubing Sun and colleagues at UMass Amherst.

**Methods**

**Encapsulation Efficiency**

Encapsulation efficiency (EE) of HLNP formulations was determined using the Quant-iT™ RiboGreen RNA assay (Thermo Fisher Scientific) with separate standard curves established under matched lysis conditions. HLNP samples were diluted in TE buffer to fall within the linear detection range. Free (unencapsulated) RNA was quantified by incubating diluted HLNPs with RiboGreen reagent in the absence of detergent. Total RNA was quantified in parallel samples following lysis with 2% Triton X-100 for 10 min at room temperature to ensure complete release of encapsulated RNA. Standard curves were generated using the same cargo RNA under matched buffer conditions with or without 2% Triton X-100. Fluorescence was measured after 5 min incubation at room temperature in the dark (Ex 485 nm, Em 528 nm). Encapsulation efficiency (EE%) was calculated as:

EE% = [(Total RNA − Free RNA) / Total RNA] × 100%

**Cell Binding Analysis**

bEnd.3 cells were seeded into 96-well plates and allowed to adhere overnight. HLNPs were pre-incubated with ApoA-I or vitronectin at a mass ratio of 10:1 (HLNP : protein) for 2 h at 37 °C to allow protein corona formation. HLNPs encapsulating Cy5-labeled DNA (200 nM) were then added to the cells and incubated at 4 °C for 1 h in Opti-MEM to permit membrane binding while minimizing endocytosis. Cells were subsequently washed with PBS to remove unbound nanoparticles and stained with Hoechst 33342 (1 μg mL^-1^) for 15 min. Cell-associated fluorescence was quantified using a multimode plate reader (Spark, Tecan), and nanoparticle binding to the cell membrane was visualized by confocal laser scanning microscopy (AXR-NSPARC, Nikon).

**Endocytic Pathway Analysis of HLNP Uptake**

To investigate the endocytic pathways involved in HLNP uptake, pharmacological inhibition assays were performed in bEnd.3 cells. Cells were seeded in 96-well plates at a density of 1 × 10^4^ cells per well and cultured to ~70% confluence. Prior to nanoparticle treatment, cells were pre-incubated for 30 min at 37 °C with specific endocytic inhibitors: chlorpromazine (clathrin-mediated endocytosis), genistein (caveolae-mediated endocytosis), methyl-β-cyclodextrin (cholesterol depletion), dynasore (dynamin-dependent endocytosis), EIPA (macropinocytosis), or cytochalasin D (actin polymerization). Cells were then treated with HLNPs encapsulating luciferase mRNA, which were pre-incubated with 10% (v/v) mouse serum at 37 °C for 1 h to allow protein corona formation, at a fixed mRNA dose in the continued presence of inhibitors. After 4 h, the medium was replaced with fresh complete medium without inhibitors, and cells were incubated for an additional 20 h to allow protein expression. Luciferase activity was quantified using a commercial assay kit and normalized to total cellular protein.

**HLNP penetration in brain organoids**

Brain organoids were used to evaluate the penetration and distribution of HLNP formulations within three-dimensional neural tissue. Organoids were cultured in low-attachment plates and maintained in organoid culture medium under standard conditions (37 °C, 5% CO₂). HLNPs encapsulating Cy5-labeled mRNA were added to the culture medium at the indicated concentrations and incubated with the organoids for 24 h. After incubation, organoids were gently washed three times with phosphate-buffered saline (PBS) to remove unbound nanoparticles. Organoids were then transferred to glass-bottom dishes and imaged using confocal microscopy. Three-dimensional fluorescence images were reconstructed from z-stack scans to visualize the spatial distribution and penetration of Cy5-labeled ssDNA delivered by different HLNP formulations.

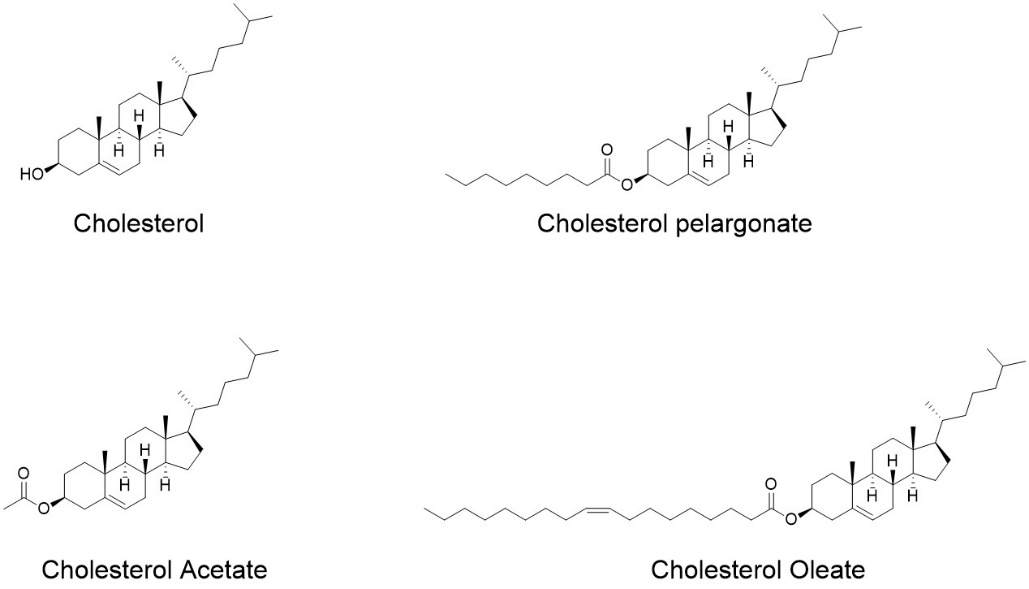

**Supplementary Figure 1**: Chemical structures of cholesterol and its ester derivatives used in this study, including cholesterol pelargonate, cholesterol acetate, and cholesterol oleate.

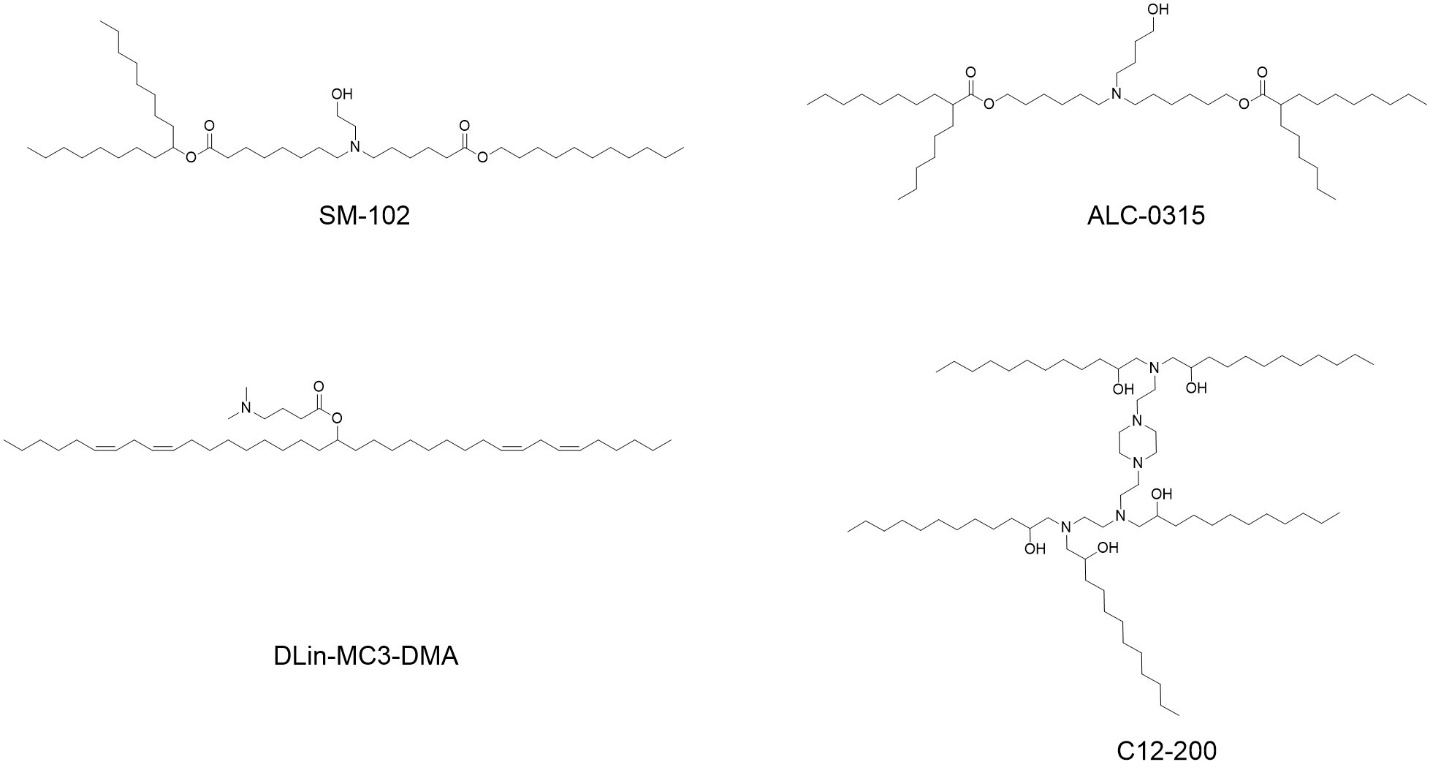

**Supplementary Figure 2**: Chemical structures of representative ionizable lipids employed in this study, including SM-102, ALC-0315, DLin-MC3-DMA, and C12-200.

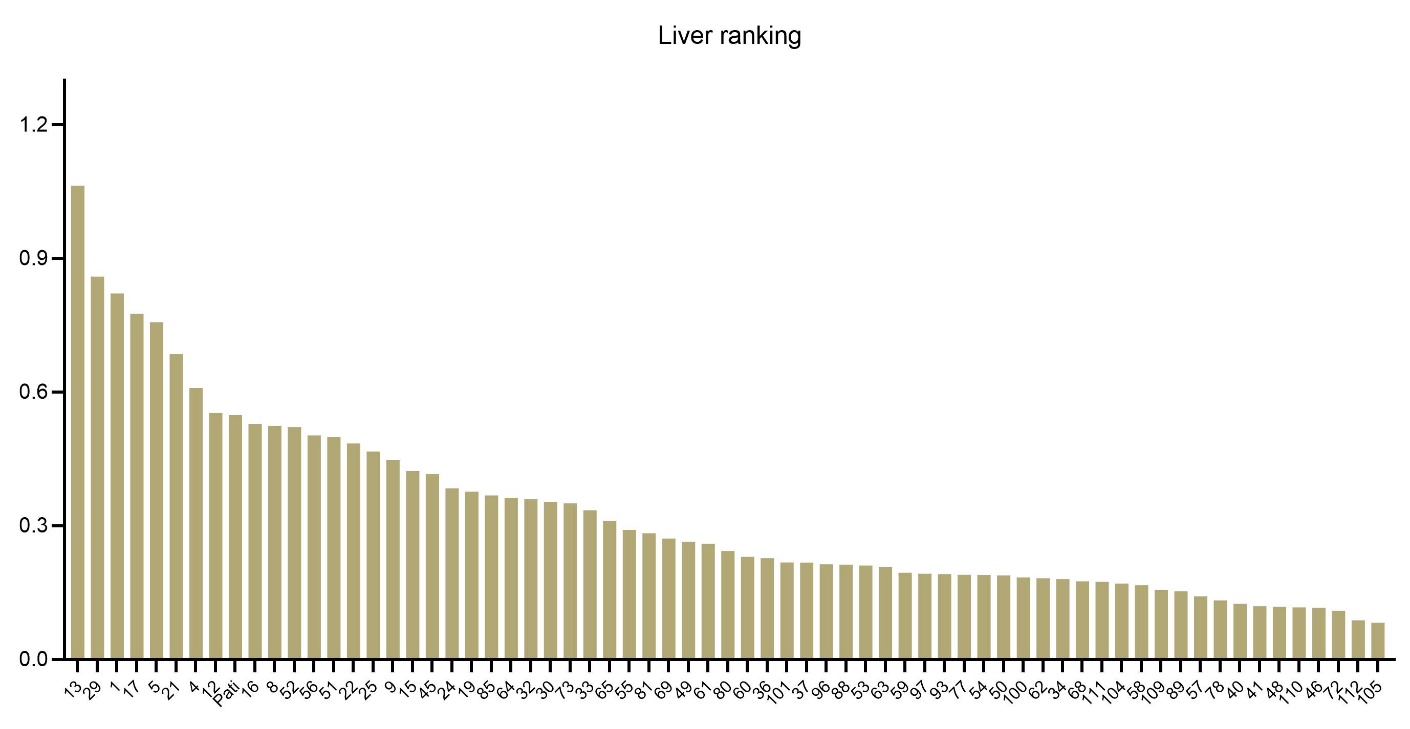

**Supplementary Figure 3:** Ranking of HLNPs based on liver accumulation following intravenous administration.

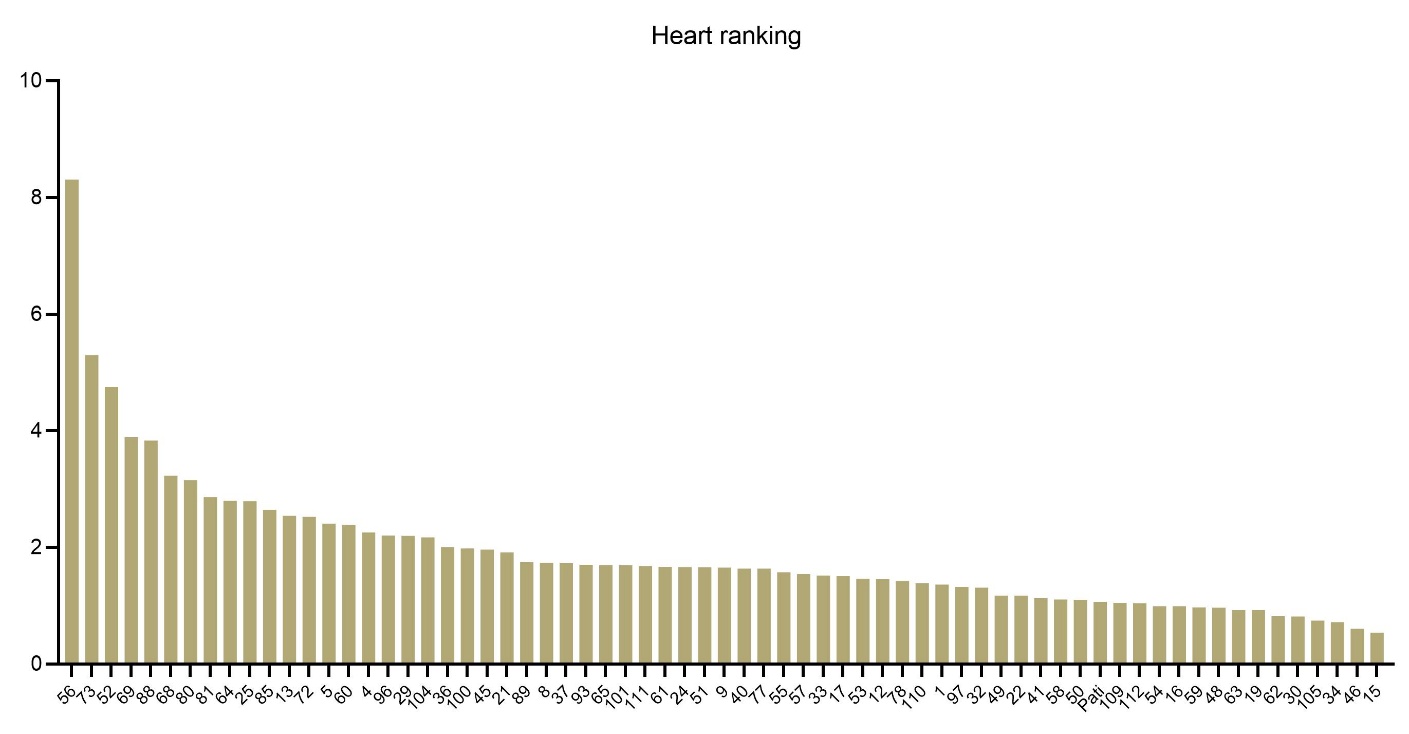

**Supplementary Figure 4:** Ranking of HLNPs based on heart accumulation following intravenous administration.

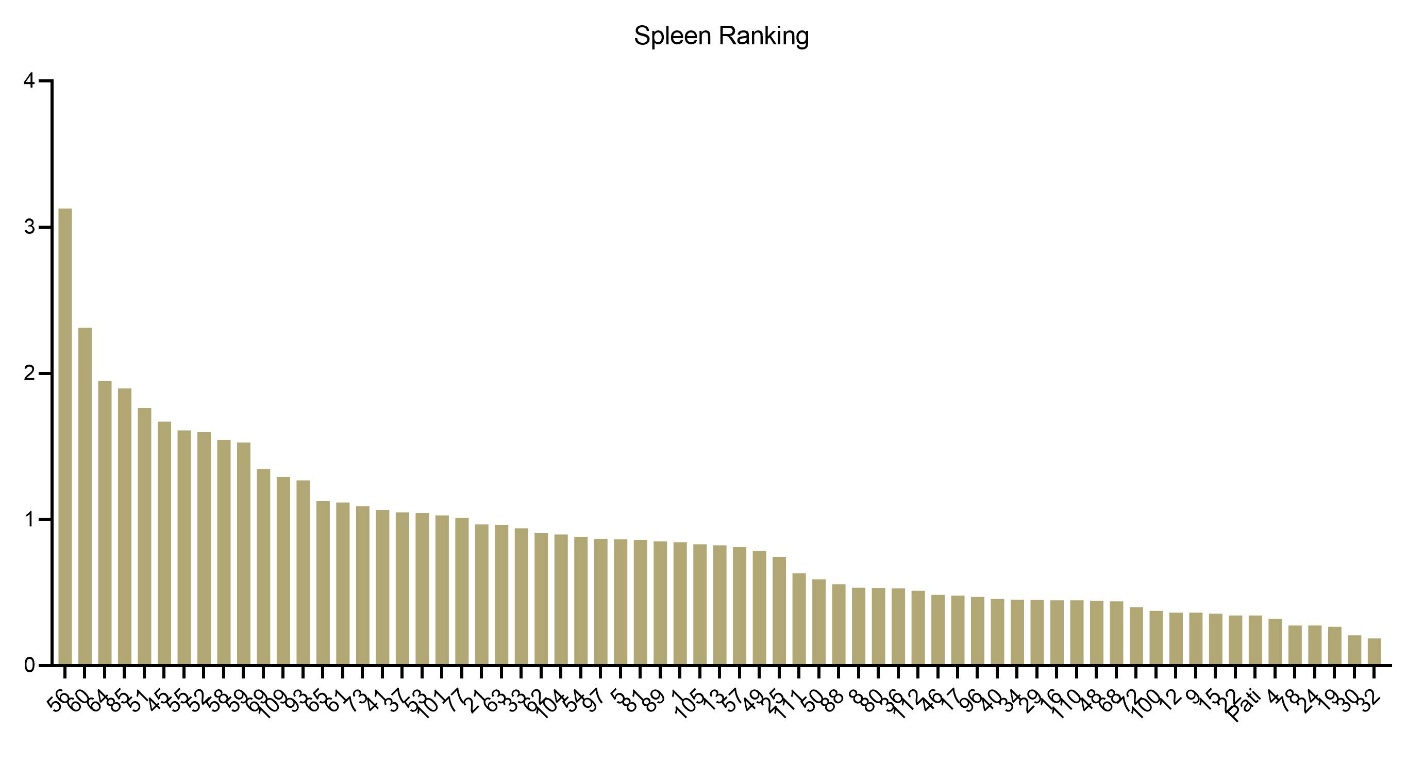

**Supplementary Figure 5:** Ranking of HLNPs based on spleen accumulation following intravenous administration.

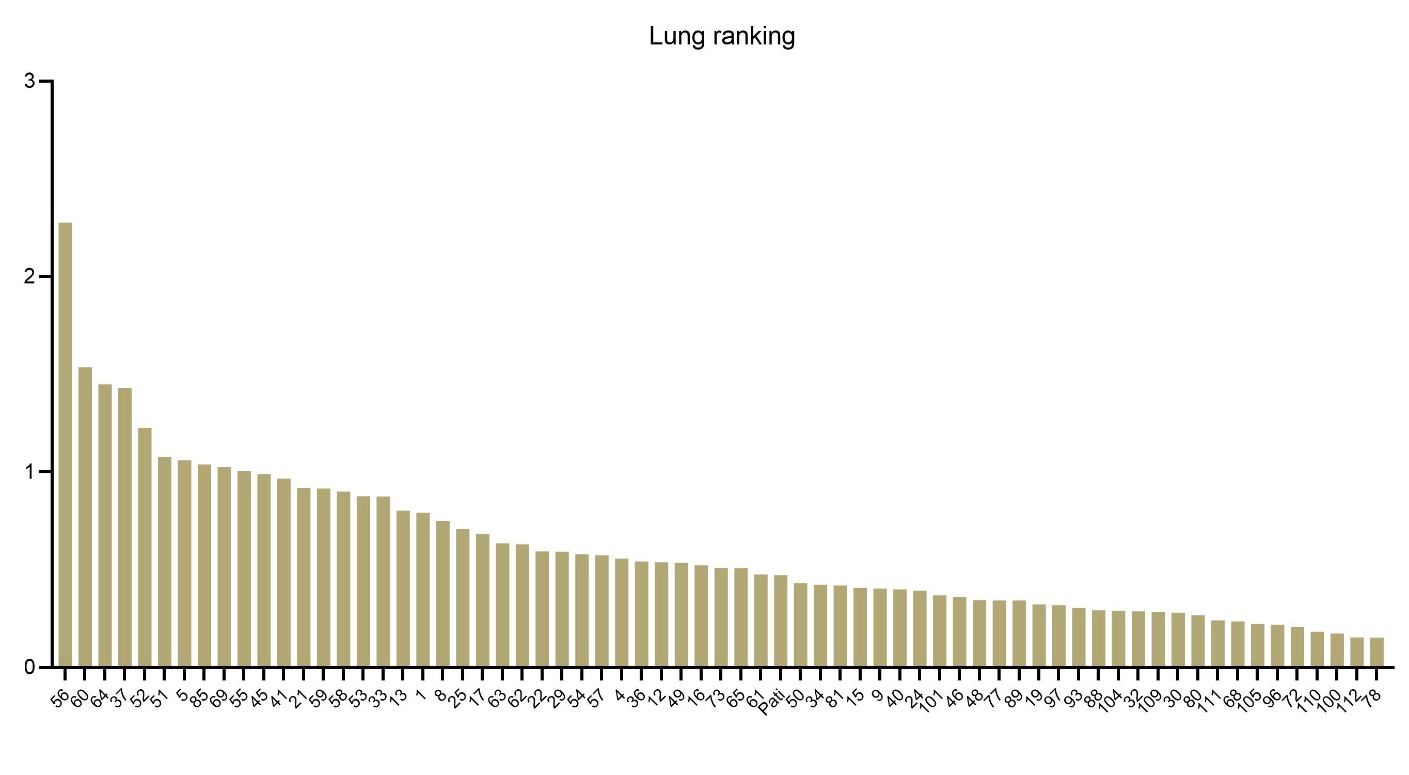
**Supplementary Figure 6:** Ranking of HLNPs based on lung accumulation following intravenous administration.

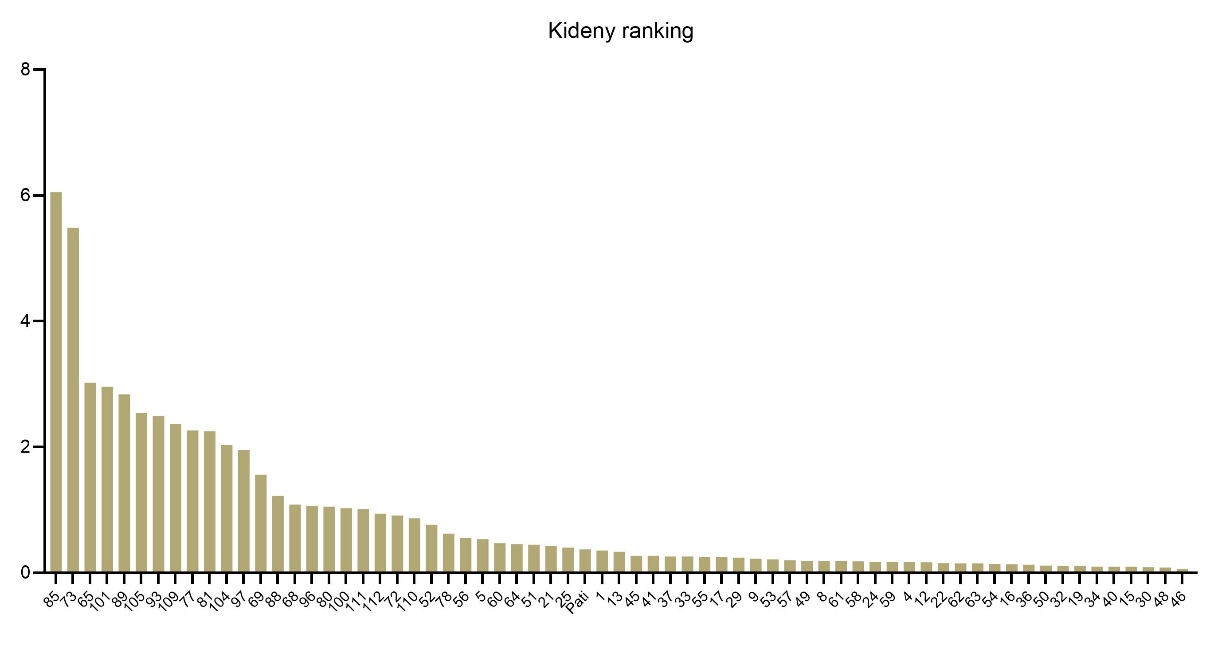

**Supplementary Figure 7:** Ranking of HLNPs based on kidney accumulation following intravenous administration.

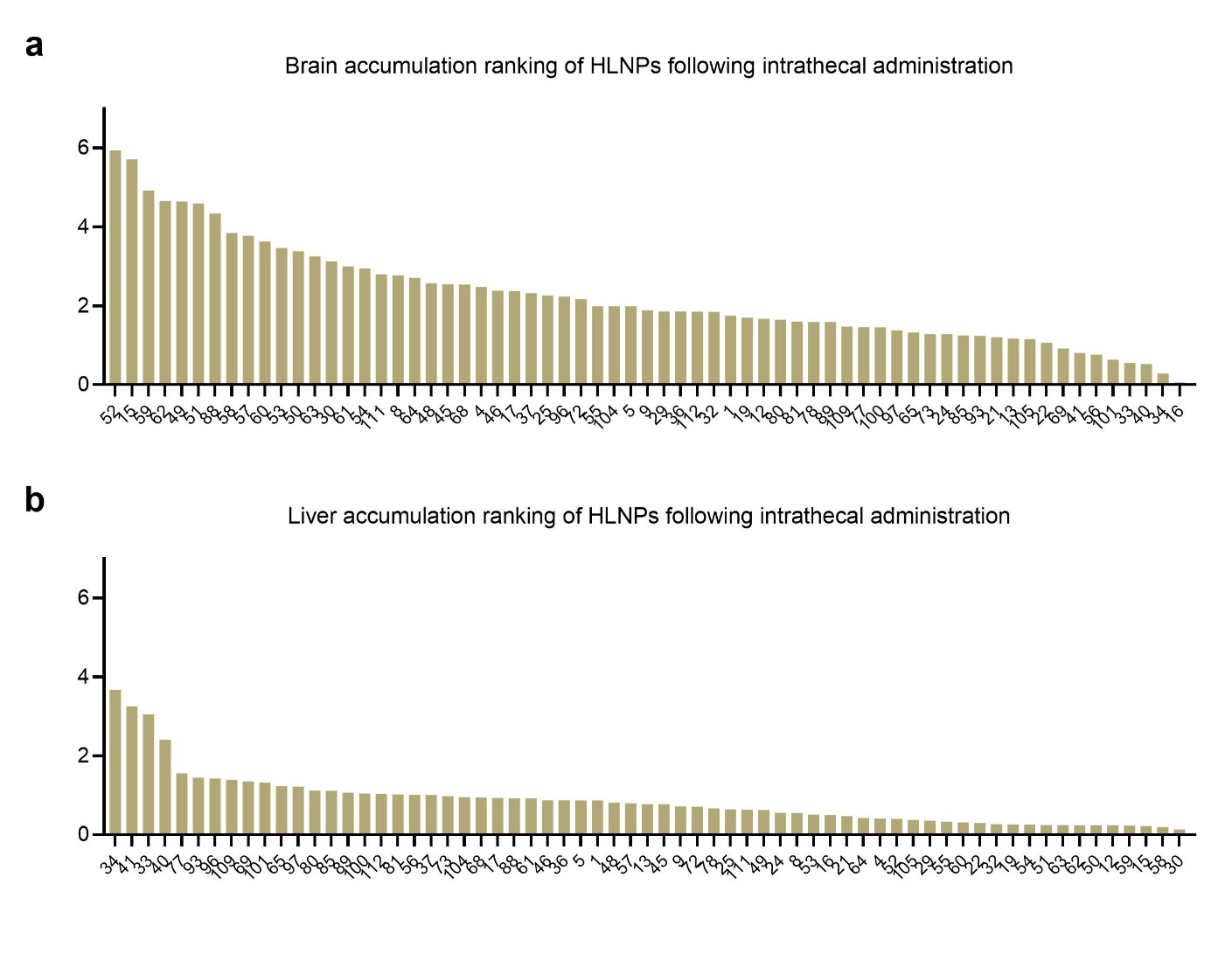
**Supplementary Figure 8：** Tissue-specific ranking of HLNPs after intrathecal delivery. **a**, Brain accumulation ranking. **b**, Liver accumulation ranking.

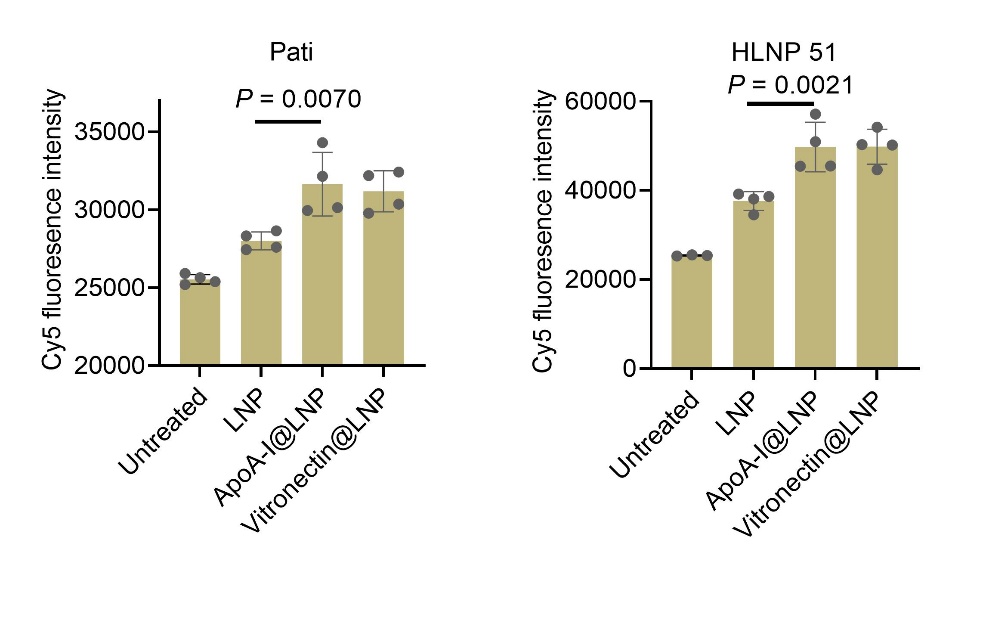

**Supplementary Figure 9:** Cy5 fluorescence intensity in bEnd.3 cells following incubation with Cy5 encapsulated HLNPs at 4 °C were quantified by plate reader to assess nanoparticle surface association while minimizing endocytosis. Compared with untreated controls, HLNP-treated cells showed increased Cy5 signal. Pre-coating with ApoA-I or vitronectin further enhanced nanoparticle association, consistent with fluorescence imaging results. Data are presented as mean ± s.d. Statistical significance was determined by one-way ANOVA with appropriate post hoc tests. *n* = 4 for all groups, except for the untreated group in HLNP 51, where *n* = 3.

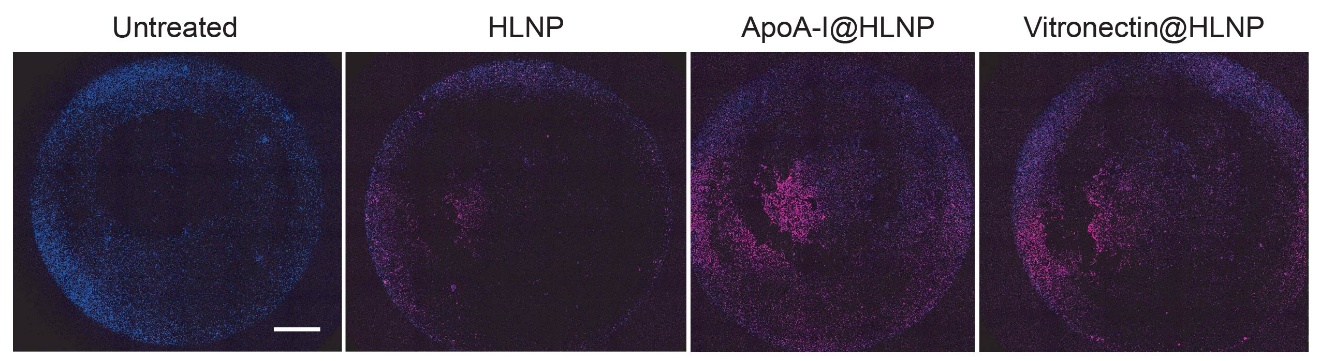

**Supplementary Figure 10:** Enhanced binding of Cy5-encapsulated HLNP 51 to bEnd.3 cells at 4 °C. Fluorescence images showing surface binding of HLNP to bEnd.3 cells at 4 °C. ApoA-I– and Vitronectin-coated nanoparticles exhibited increased association compared to uncoated HLNPs, indicating protein corona–mediated enhancement of endothelial binding. The scale bar is 1000 μm.

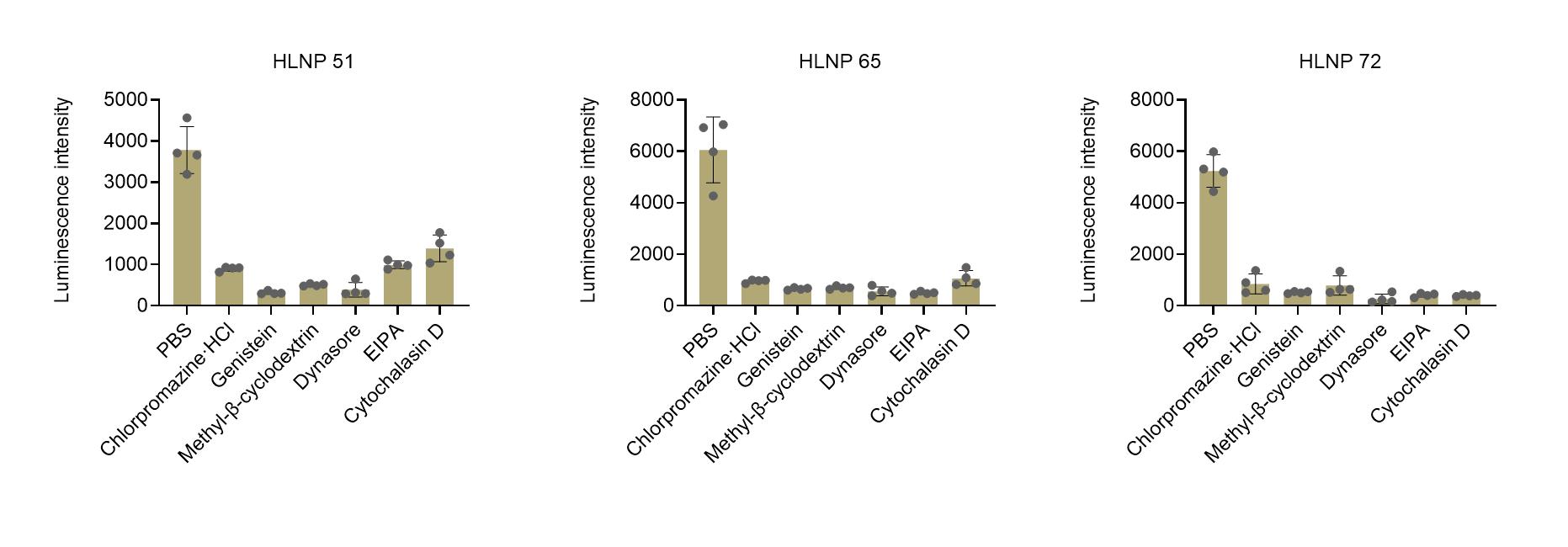

**Supplementary Figure 11:** Endocytic pathway analysis of HLNP uptake in bEnd.3 cells. Cells were pre-treated with pharmacological inhibitors targeting distinct endocytic pathways, including chlorpromazine (10 µg mL⁻¹, clathrin-mediated endocytosis), genistein (200 µM, caveolae-mediated uptake), methyl-β-cyclodextrin (5 mM, cholesterol depletion), dynasore (80 µM, dynamin-dependent endocytosis), EIPA (50 µM, macropinocytosis), and cytochalasin D (5 µM, actin polymerization). Cells were then incubated with HLNPs encapsulating luciferase mRNA that had been pre-conditioned with 10% (v/v) mouse serum to allow protein corona formation. Luciferase activity was measured after 24 h and normalized to total cellular protein. Data are presented as mean ± s.d. (*n* = 3).

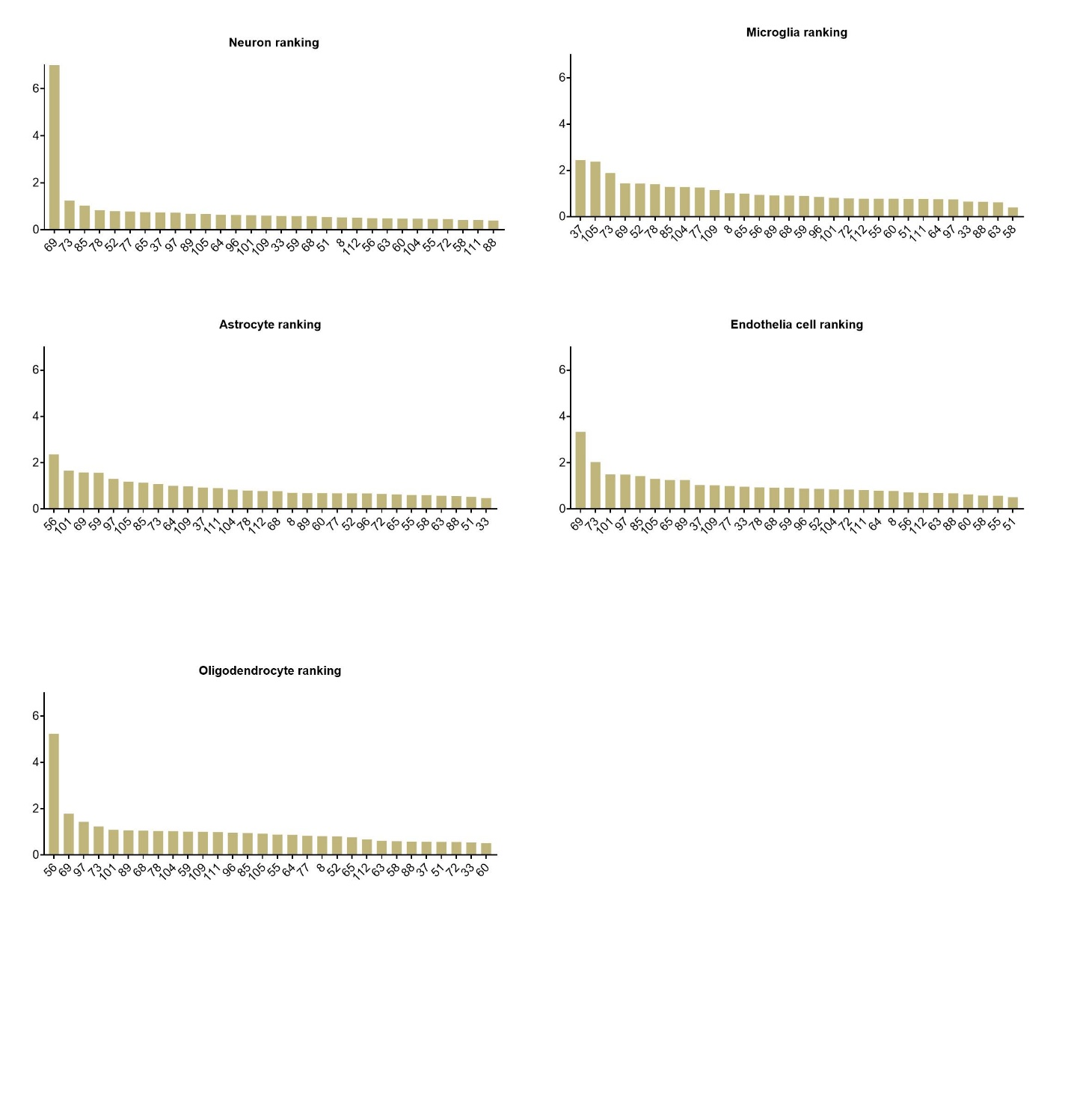

**Supplementary Figure 12**: Cell-type–specific enrichment of HLNP formulations in the brain. Bar plots showing the ranking of HLNP formulations based on normalized barcode enrichment within neurons, microglia, astrocytes, endothelial cells, and oligodendrocytes following in vivo pooled screening.

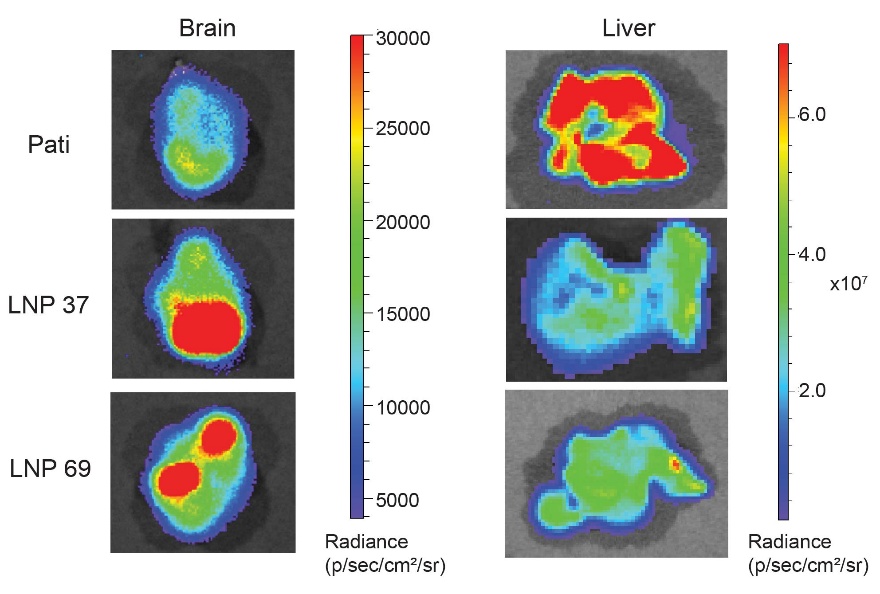

**Supplementary Figure 13:** Representative ex vivo bioluminescence images of mouse brains and major organs following systemic administration of FLuc mRNA–loaded HLNP formulations (Pati, HLNP 37 and HLNP 69), illustrating differences in biodistribution.

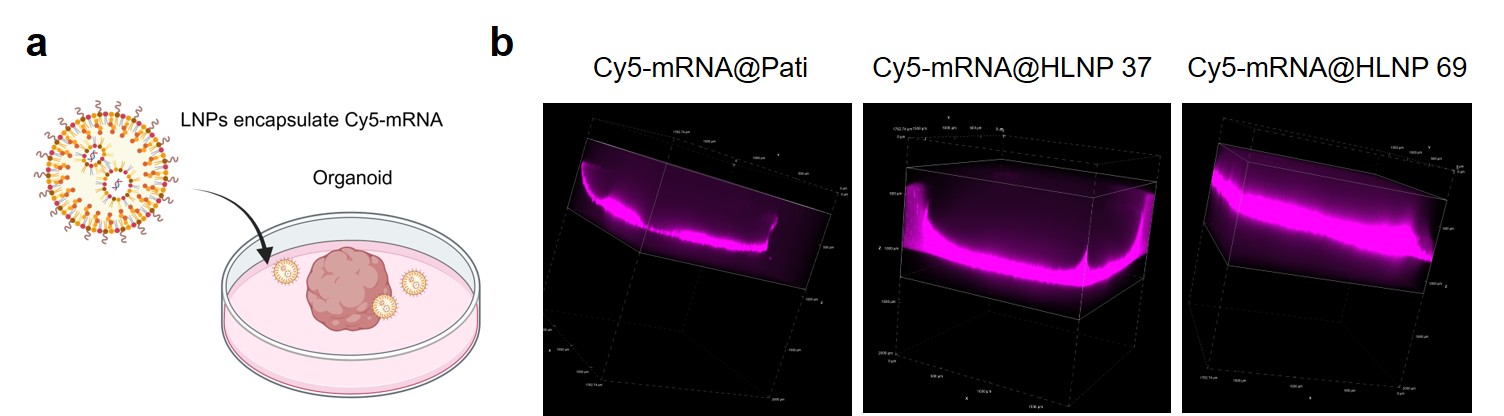

**Supplementary Figure 14:** Delivery of Cy5-mRNA HLNPs in brain organoids.
**a**, Schematic illustration of the experimental design. HLNPs encapsulating Cy5-labeled mRNA were incubated with brain organoids to evaluate nanoparticle penetration and distribution within the three-dimensional tissue structure. **b**, Representative three-dimensional fluorescence reconstructions showing the distribution of Cy5 signal in organoids treated with Pati-Cy5, 37-Cy5, and 69-Cy5 HLNP formulations. Fluorescence intensity corresponds to the presence and penetration of Cy5-labeled mRNA delivered by the respective HLNPs.

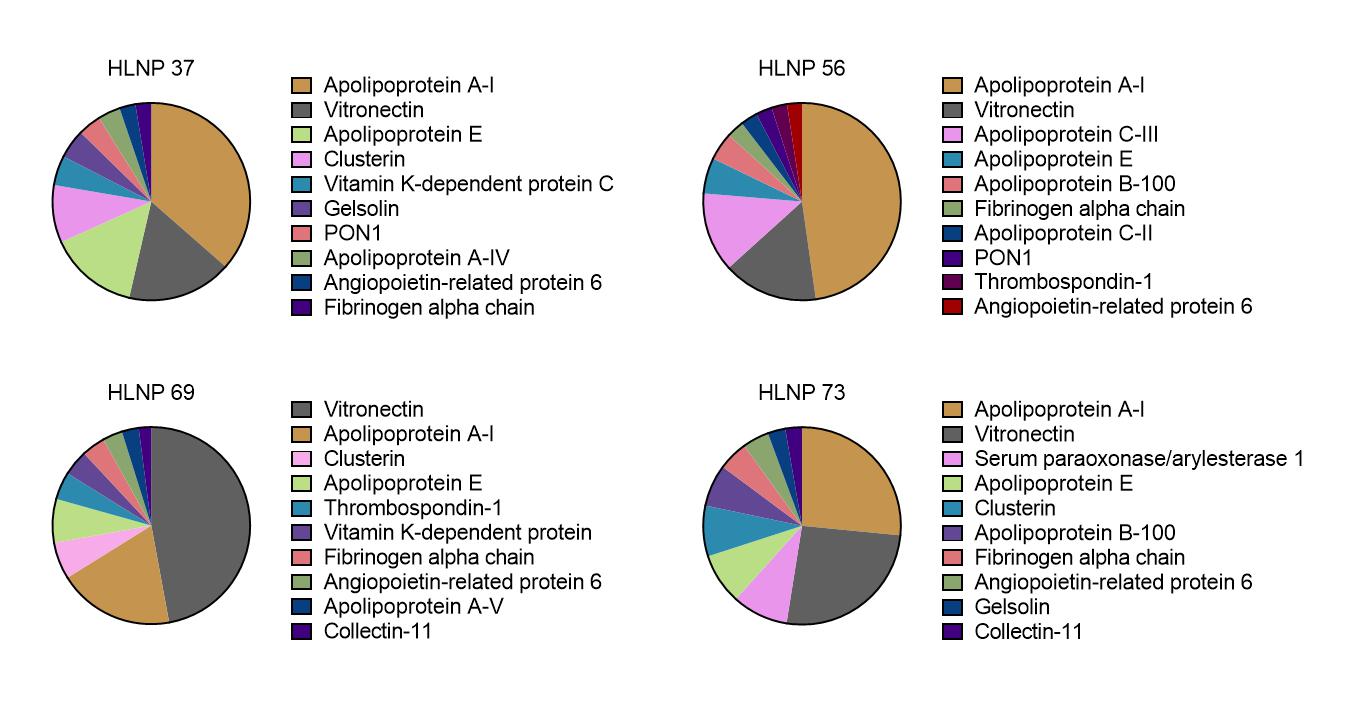

**Supplementary Figure 15**: Comparative analysis of protein corona composition associated with HLNP formulations. Pie charts showing the relative abundance of the top 10 serum proteins identified in the protein corona formed on representative HLNP formulations (37, 56, 69, and 73) following incubation with mouse serum. Major corona constituents include apolipoprotein A-I (ApoA-I), vitronectin, and apolipoprotein E (ApoE), along with other serum proteins. The relative enrichment of ApoA-I, vitronectin, and ApoE among the top-ranked corona proteins is highlighted for each formulation.

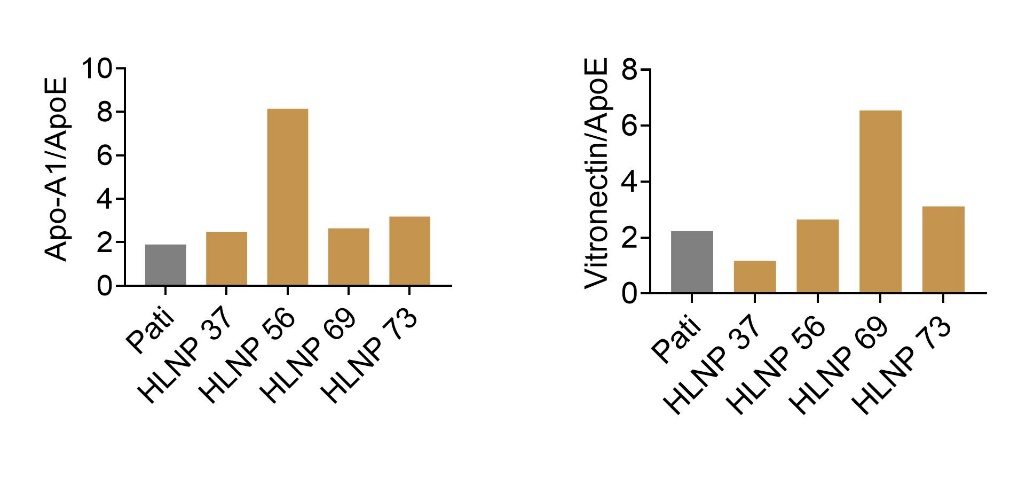

**Supplementary Figure 16:** Ratio-based analysis of protein corona composition across HLNP formulations. Bar plots showing the ratios of ApoA-I/ApoE and vitronectin/ApoE for representative HLNP formulations relative to the Pati control.

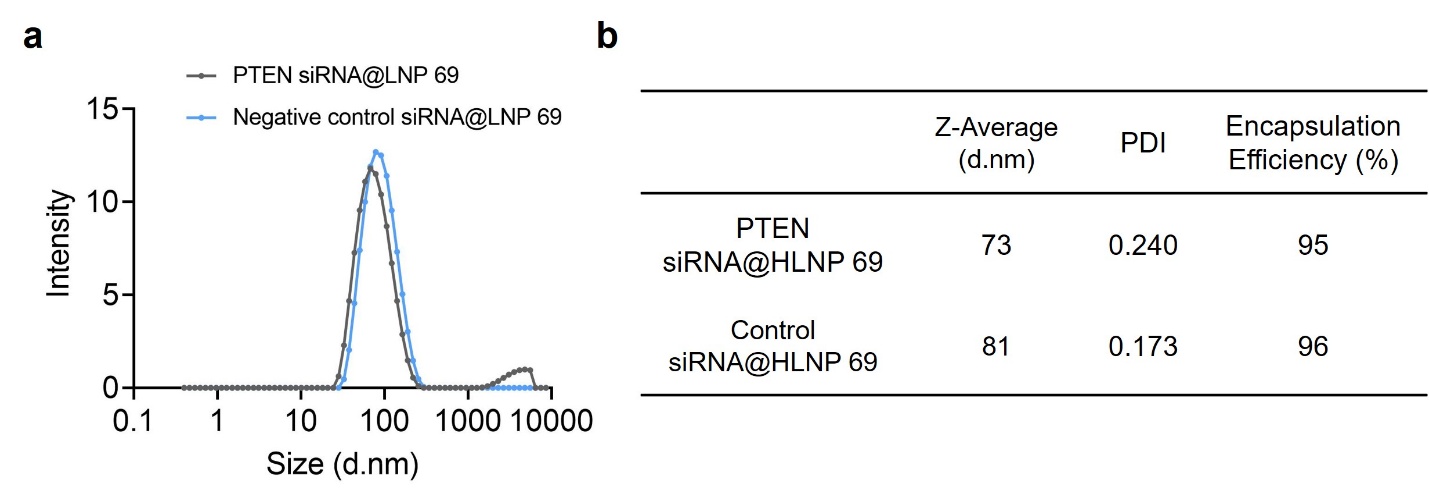

**Supplementary Figure 17:** Physicochemical characterization of siRNA-loaded HLNP 69 formulations. **a**, Hydrodynamic size distribution profiles of PTEN siRNA@HLNP 69 and control siRNA@HLNP 69 measured by dynamic light scattering (DLS). **b**, Summary of physicochemical properties, including Z-average diameter, polydispersity index (PDI), and encapsulation efficiency (EE).

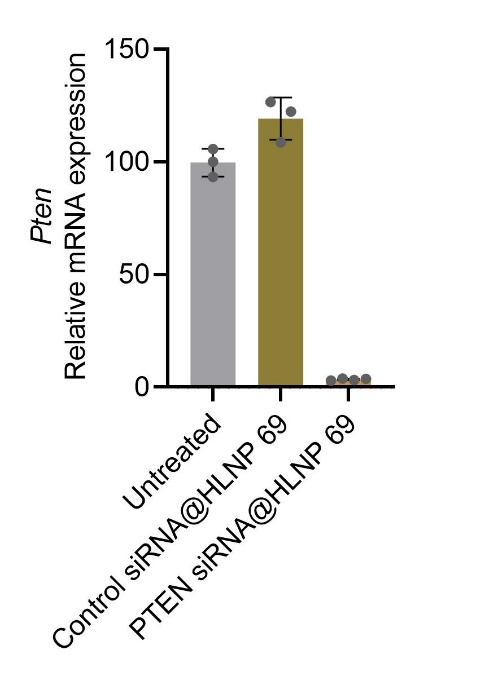

**Supplementary Figure 18:** In vitro validation of PTEN knockdown in N2a cells following treatment with HLNP 69 encapsulating PTEN siRNA, as quantified by RT–qPCR (n = 3 for untreated, n = 4 for control siRNA, and n = 4 for PTEN siRNA).

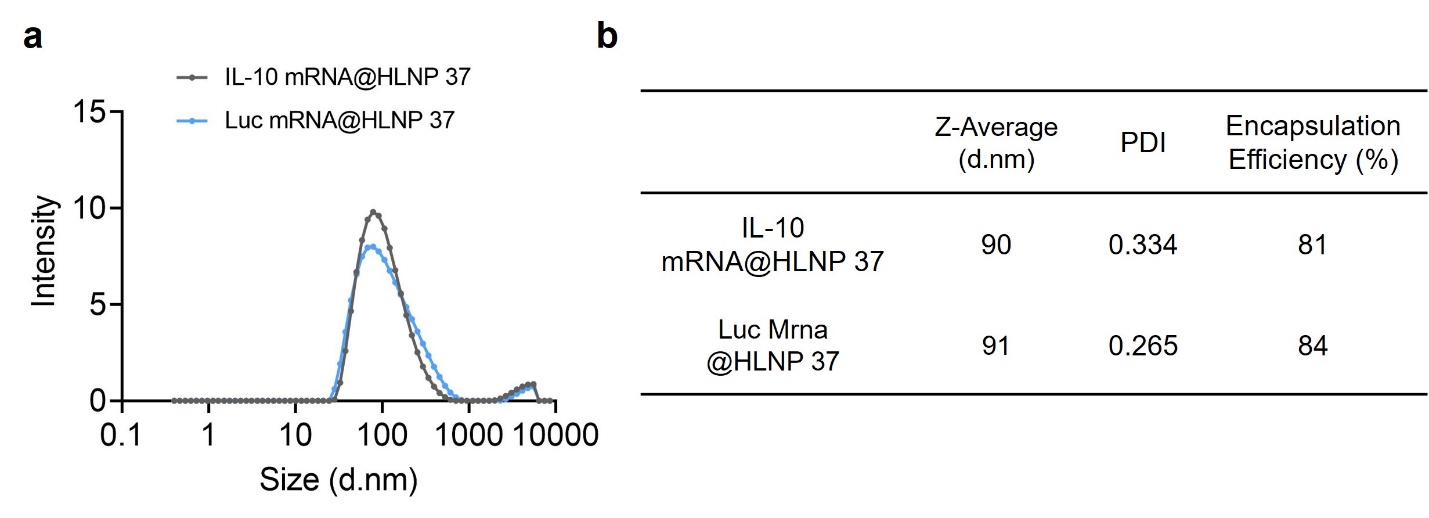

**Supplementary Figure 19:** Physicochemical characterization of mRNA-loaded HLNP 37 formulations. **a**, Hydrodynamic size distribution profiles of PTEN siRNA@HLNP 37 and control siRNA@HLNP 37 measured by dynamic light scattering (DLS). **b**, Summary of physicochemical properties, including Z-average diameter, polydispersity index (PDI), and encapsulation efficiency.

**Supplementary Table 1: Physicochemical characterization of LNPs, including size, polydispersity index (PDI), and encapsulation efficiency**

|  | **LNP Components**  **Help lipid (10) : Cholesterol (38.5) : Ionizable lipids (50%): DMG-PEG2000 (1.5)** | **Size**  **Z-Average**  **(d.nm)** | **PDI** | **Encapsulation Efficiency (%)** |
| --- | --- | --- | --- | --- |
| 1 | 16:0-18:2 PC/Cholesterol/SM-102/DMG-PEG2000 | 95 | 0.237 | 84 |
| 2 | 16:0-18:2 PC/Cholesterol pelargonate/SM-102/DMG-PEG2000 | 212 | 0.445 | 37 |
| 3 | 16:0-18:2 PC/Cholesterol oleate/SM-102/DMG-PEG2000 | 132 | 0.217 | 44 |
| 4 | 16:0-18:2 PC/Cholesterol acetate/SM-102/DMG-PEG2000 | 159 | 0.284 | 72 |
| 5 | 16:0-18:2 PC/Cholesterol/Dlin-MC3-DMA/DMG-PEG2000 | 175 | 0.11 | 84 |
| 6 | 16:0-18:2 PC/Cholesterol pelargonate/Dlin-MC3-DMA/DMG-PEG2000 | 197 | 0.189 | 44 |
| 7 | 16:0-18:2 PC/Cholesterol oleate/Dlin-MC3-DMA/DMG-PEG2000 | 117 | 0.246 | 41 |
| 8 | 16:0-18:2 PC/Cholesterol acetate /Dlin-MC3-DMA/DMG-PEG2000 | 159 | 0.186 | 75 |
| 9 | 16:0-18:2 PC/Cholesterol/ALC-0315/DMG-PEG2000 | 148 | 0.311 | 73 |
| 10 | 16:0-18:2 PC/Cholesterol pelargonate/ALC-0315/DMG-PEG2000 | 200 | 0.44 | 19 |
| 11 | 16:0-18:2 PC/Cholesterol oleate/ALC-0315/DMG-PEG2000 | 187 | 0.455 | 45 |
| 12 | 16:0-18:2 PC/Cholesterol acetate/ALC-0315/DMG-PEG2000 | 168 | 0.365 | 72 |
| 13 | 16:0-18:2 PC/Cholesterol/C12-200/DMG-PEG2000 | 134 | 0.209 | 83 |
| 14 | 16:0-18:2 PC/Cholesterol pelargonate / C12-200/DMG-PEG2000 | 111 | 0.208 | 34 |
| 15 | 16:0-18:2 PC/Cholesterol oleate/C12-200/DMG-PEG2000 | 165 | 0.267 | 75 |
| 16 | 16:0-18:2 PC/Cholesterol acetate/C12-200/DMG-PEG2000 | 149 | 0.271 | 77 |
| 17 | 18:0-18:2 PC/Cholesterol/SM-102/DMG-PEG2000 | 128 | 0.243 | 90 |
| 18 | 18:0-18:2 PC/Cholesterol pelargonate/SM-102/DMG-PEG2000 | 111 | 0.323 | 56 |
| 19 | 18:0-18:2 PC/Cholesterol oleate/SM-102/DMG-PEG2000 | 116 | 0.339 | 74 |
| 20 | 18:0-18:2 PC/Cholesterol acetate/SM-102/DMG-PEG2000 | 112 | 0.318 | 46 |
| 21 | 18:0-18:2 PC/Cholesterol/Dlin-MC3-DMA/DMG-PEG2000 | 156 | 0.303 | 87 |
| 22 | 18:0-18:2 PC/Cholesterol pelargonate/Dlin-MC3-DMA/DMG-PEG2000 | 156 | 0.415 | 71 |
| 23 | 18:0-18:2 PC/Cholesterol oleate/Dlin-MC3-DMA/DMG-PEG2000 | 138 | 0.341 | 39 |
| 24 | 18:0-18:2 PC/Cholesterol acetate/Dlin-MC3-DMA/DMG-PEG2000 | 106 | 0.372 | 71 |
| 25 | 18:0-18:2 PC/Cholesterol/ALC-0315/DMG-PEG2000 | 107 | 0.326 | 84 |
| 26 | 18:0-18:2 PC/Cholesterol pelargonate/ALC-0315/DMG-PEG2000 | 116 | 0.343 | 41 |
| 27 | 18:0-18:2 PC/Cholesterol oleate/ALC-0315/DMG-PEG2000 | 124 | 0.248 | 27 |
| 28 | 18:0-18:2 PC/Cholesterol acetate/ALC-0315/DMG-PEG2000 | 110 | 0.291 | 44 |
| 29 | 18:0-18:2 PC/Cholesterol/C12-200/DMG-PEG2000 | 109 | 0.217 | 85 |
| 30 | 18:0-18:2 PC/Cholesterol pelargonate/C12-200/DMG-PEG2000 | 113 | 0.221 | 71 |
| 31 | 18:0-18:2 PC/Cholesterol oleate/C12-200/DMG-PEG2000 | 173 | 0.283 | 46 |
| 32 | 18:0-18:2 PC/Cholesterol acetate/C12-200/DMG-PEG2000 | 117 | 0.248 | 75 |
| 33 | 16:0-18:1 PC/Cholesterol/SM-102/DMG-PEG2000 | 126 | 0.223 | 86 |
| 34 | 16:0-18:1 PC/Cholesterol pelargonate /SM-102/DMG-PEG2000 | 129 | 0.236 | 70 |
| 35 | 16:0-18:1 PC/Cholesterol oleate /SM-102/DMG-PEG2000 | 140 | 0.255 | 44 |
| 36 | 16:0-18:1 PC/Cholesterol acetate /SM-102/DMG-PEG2000 | 144 | 0.246 | 74 |
| 37 | 16:0-18:1 PC/Cholesterol/Dlin-MC3-DMA/DMG-PEG2000 | 193 | 0.255 | 86 |
| 38 | 16:0-18:1 PC/Cholesterol pelargonate /Dlin-MC3-DMA/DMG-PEG2000 | 161 | 0.247 | 37 |
| 39 | 16:0-18:1 PC/Cholesterol oleate /Dlin-MC3-DMA/DMG-PEG2000 | 128 | 0.239 | 37 |
| 40 | 16:0-18:1 PC/Cholesterol acetate /Dlin-MC3-DMA/DMG-PEG2000 | 142 | 0.301 | 72 |
| 41 | 16:0-18:1 PC/Cholesterol/ALC-0315/DMG-PEG2000 | 131 | 0.286 | 79 |
| 42 | 16:0-18:1 PC/Cholesterol pelargonate/ALC-0315/DMG-PEG2000 | 144 | 0.258 | 34 |
| 43 | 16:0-18:1 PC/Cholesterol oleate/ALC-0315/DMG-PEG2000 | 133 | 0.23 | 23 |
| 44 | 16:0-18:1 PC/Cholesterol acetate/ALC-0315/DMG-PEG2000 | 131 | 0.244 | 38 |
| 45 | 16:0-18:1 PC/Cholesterol/C12-200/DMG-PEG2000 | 122 | 0.221 | 90 |
| 46 | 16:0-18:1 PC/Cholesterol pelargonate/C12-200/DMG-PEG2000 | 134 | 0.218 | 65 |
| 47 | 16:0-18:1 PC/Cholesterol oleate/C12-200/DMG-PEG2000 | 123 | 0.2 | 45 |
| 48 | 16:0-18:1 PC/Cholesterol acetate/C12-200/DMG-PEG2000 | 119 | 0.198 | 70 |
| 49 | 16:0-20:4 PC/Cholesterol/SM-102/DMG-PEG2000 | 118 | 0.26 | 85 |
| 50 | 16:0-20:4 PC/Cholesterol pelargonate/SM-102/DMG-PEG2000 | 122 | 0.251 | 70 |
| 51 | 16:0-20:4 PC/Cholesterol oleate/SM-102/DMG-PEG2000 | 114 | 0.269 | 83 |
| 52 | 16:0-20:4 PC/Cholesterol acetate/SM-102/DMG-PEG2000 | 106 | 0.322 | 87 |
| 53 | 16:0-20:4 PC/Cholesterol/ Dlin-MC3-DMA /DMG-PEG2000 | 125 | 0.325 | 87 |
| 54 | 16:0-20:4 PC/Cholesterol pelargonate/Dlin-MC3-DMA /DMG-PEG2000 | 149 | 0.23 | 75 |
| 55 | 16:0-20:4 PC/Cholesterol oleate/Dlin-MC3-DMA /DMG-PEG2000 | 148 | 0.379 | 75 |
| 56 | 16:0-20:4 PC/Cholesterol acetate/Dlin-MC3-DMA /DMG-PEG2000 | 142 | 0.458 | 88 |
| 57 | 16:0-20:4 PC/Cholesterol/ALC-0315/DMG-PEG2000 | 125 | 0.234 | 88 |
| 58 | 16:0-20:4 PC/Cholesterol pelargonate/ALC-0315/DMG-PEG2000 | 120 | 0.266 | 83 |
| 59 | 16:0-20:4 PC/Cholesterol oleate/ALC-0315/DMG-PEG2000 | 114 | 0.244 | 85 |
| 60 | 16:0-20:4 PC/Cholesterol acetate/ALC-0315/DMG-PEG2000 | 111 | 0.367 | 91 |
| 61 | 16:0-20:4 PC/Cholesterol/C12-200/DMG-PEG2000 | 98 | 0.255 | 92 |
| 62 | 16:0-20:4 PC/Cholesterol pelargonate/C12-200/DMG-PEG2000 | 181 | 0.122 | 76 |
| 63 | 16:0-20:4 PC/Cholesterol oleate/C12-200/DMG-PEG2000 | 196 | 0.12 | 74 |
| 64 | 16:0-20:4 PC/Cholesterol acetate/C12-200/DMG-PEG2000 | 107 | 0.273 | 91 |
| 65 | 16:0 SM/Cholesterol/SM-102/DMG-PEG2000 | 120 | 0.26 | 77 |
| 66 | 16:0 SM/Cholesterol pelargonate/SM-102/DMG-PEG2000 | 100 | 0.313 | 38 |
| 67 | 16:0 SM/Cholesterol oleate/SM-102/DMG-PEG2000 | 103 | 0.287 | 31 |
| 68 | 16:0 SM/Cholesterol acetate/SM-102/DMG-PEG2000 | 127 | 0.268 | 72 |
| 69 | 16:0 SM/Cholesterol/ Dlin-MC3-DMA /DMG-PEG2000 | 171 | 0.242 | 77 |
| 70 | 16:0 SM/Cholesterol pelargonate/Dlin-MC3-DMA /DMG-PEG2000 | 107 | 0.346 | 25 |
| 71 | 16:0 SM/Cholesterol oleate/Dlin-MC3-DMA /DMG-PEG2000 | 98 | 0.311 | 26 |
| 72 | 16:0 SM/Cholesterol acetate/Dlin-MC3-DMA /DMG-PEG2000 | 121 | 0.34 | 74 |
| 73 | 16:0 SM/Cholesterol/ ALC-0315/DMG-PEG2000 | 122 | 0.196 | 81 |
| 74 | 16:0 SM/Cholesterol pelargonate/ALC-0315/DMG-PEG2000 | 96 | 0.282 | 46 |
| 75 | 16:0 SM/Cholesterol oleate/ALC-0315/DMG-PEG2000 | 95 | 0.279 | 41 |
| 76 | 16:0 SM/Cholesterol acetate/ALC-0315/DMG-PEG2000 | 97 | 0.276 | 37 |
| 77 | 16:0 SM/Cholesterol/C12-200/DMG-PEG2000 | 101 | 0.3 | 81 |
| 78 | 16:0 SM/Cholesterol pelargonate/C12-200/DMG-PEG2000 | 92 | 0.224 | 76 |
| 79 | 16:0 SM/Cholesterol oleate/C12-200/DMG-PEG2000 | 88 | 0.163 | 49 |
| 80 | 16:0 SM/Cholesterol acetate/C12-200/DMG-PEG2000 | 94 | 0.187 | 74 |
| 81 | 18:0 SM/Cholesterol/SM-102/DMG-PEG2000 | 97 | 0.275 | 77 |
| 82 | 18:0 SM/Cholesterol pelargonate/SM-102/DMG-PEG2000 | 119 | 0.272 | 32 |
| 83 | 18:0 SM/Cholesterol oleate/SM-102/DMG-PEG2000 | 124 | 0.288 | 22 |
| 84 | 18:0 SM/Cholesterol acetate/SM-102/DMG-PEG2000 | 122 | 0.346 | 43 |
| 85 | 18:0 SM/Cholesterol/ Dlin-MC3-DMA /DMG-PEG2000 | 90 | 0.25 | 75 |
| 86 | 18:0 SM/Cholesterol pelargonate/Dlin-MC3-DMA /DMG-PEG2000 | 122 | 0.249 | 43 |
| 87 | 18:0 SM/Cholesterol oleate/Dlin-MC3-DMA /DMG-PEG2000 | 105 | 0.268 | 21 |
| 88 | 18:0 SM/Cholesterol acetate/Dlin-MC3-DMA /DMG-PEG2000 | 121 | 0.185 | 70 |
| 89 | 18:0 SM/Cholesterol/ ALC-0315/DMG-PEG2000 | 104 | 0.251 | 79 |
| 90 | 18:0 SM/Cholesterol pelargonate/ALC-0315/DMG-PEG2000 | 104 | 0.347 | 16 |
| 91 | 18:0 SM/Cholesterol oleate/ALC-0315/DMG-PEG2000 | 110 | 0.312 | 17 |
| 92 | 18:0 SM/Cholesterol acetate/ALC-0315/DMG-PEG2000 | 108 | 0.272 | 25 |
| 93 | 18:0 SM/Cholesterol/C12-200/DMG-PEG2000 | 98 | 0.263 | 75 |
| 94 | 18:0 SM/Cholesterol pelargonate/C12-200/DMG-PEG2000 | 95 | 0.07 | 26 |
| 95 | 18:0 SM/Cholesterol oleate/C12-200/DMG-PEG2000 | 93 | 0.11 | 17 |
| 96 | 18:0 SM/Cholesterol acetate/C12-200/DMG-PEG2000 | 88 | 0.213 | 76 |
| 97 | 24:0 SM/Cholesterol/SM-102/DMG-PEG2000 | 82 | 0.186 | 79 |
| 98 | 24:0 SM/Cholesterol pelargonate/SM-102/DMG-PEG2000 | 73 | 0.265 | 25 |
| 99 | 24:0 SM/Cholesterol oleate/SM-102/DMG-PEG2000 | 86 | 0.282 | 13 |
| 100 | 24:0 SM/Cholesterol acetate/SM-102/DMG-PEG2000 | 95 | 0.278 | 72 |
| 101 | 24:0 SM/Cholesterol/ Dlin-MC3-DMA /DMG-PEG2000 | 109 | 0.322 | 84 |
| 102 | 24:0 SM/Cholesterol pelargonate/Dlin-MC3-DMA /DMG-PEG2000 | 92 | 0.213 | 17 |
| 103 | 24:0 SM/Cholesterol oleate/Dlin-MC3-DMA /DMG-PEG2000 | 82 | 0.158 | 41 |
| 104 | 24:0 SM/Cholesterol acetate/Dlin-MC3-DMA /DMG-PEG2000 | 106 | 0.207 | 73 |
| 105 | 24:0 SM/Cholesterol/ ALC-0315/DMG-PEG2000 | 84 | 0.309 | 80 |
| 106 | 24:0 SM/Cholesterol pelargonate/ALC-0315/DMG-PEG2000 | 84 | 0.211 | 45 |
| 107 | 24:0 SM/Cholesterol oleate/ALC-0315/DMG-PEG2000 | 111 | 0.357 | 43 |
| 108 | 24:0 SM/Cholesterol acetate/ALC-0315/DMG-PEG2000 | 81 | 0.309 | 17 |
| 109 | 24:0 SM/Cholesterol/C12-200/DMG-PEG2000 | 97 | 0.246 | 85 |
| 110 | 24:0 SM/Cholesterol pelargonate/C12-200/DMG-PEG2000 | 114 | 0.235 | 72 |
| 111 | 24:0 SM/Cholesterol oleate/C12-200/DMG-PEG2000 | 117 | 0.141 | 72 |
| 112 | 24:0 SM/Cholesterol acetate/C12-200/DMG-PEG2000 | 116 | 0.204 | 76 |

**Supplementary Table 2**: Barcode sequences used in this study

AG Barcode 1

5' G*A*T* GCA CGC CTT ACG ACT CAT CTN WNH TGA TAT TGN WHG TGG TTA GTC GAG CAG AGA C*T*A* G 3'

AG Barcode 2

5' G*A*T* GCA CGC CTT ACG ACT CAT CTN WNH GAC GCA ATN WHG TGG TTA GTC GAG CAG AGA C*T*A* G 3'

AG Barcode 3

5' G*A*T* GCA CGC CTT ACG ACT CAT CTN WNH GCG AGT ATN WHG TGG TTA GTC GAG CAG AGA C*T*A* G 3'

AG Barcode 4

5' G*A*T* GCA CGC CTT ACG ACT CAT CTN WNH ACC TAA TCN WHG TGG TTA GTC GAG CAG AGA C*T*A* G 3'

AG Barcode 5

5' G*A*T* GCA CGC CTT ACG ACT CAT CTN WNH AGG CGC TAN WHG TGG TTA GTC GAG CAG AGA C*T*A* G 3'

AG Barcode 6

5' G*A*T* GCA CGC CTT ACG ACT CAT CTN WNH GAT CTA CCN WHG TGG TTA GTC GAG CAG AGA C*T*A* G 3'

AG Barcode 7

5' G*A*T* GCA CGC CTT ACG ACT CAT CTN WNH CTA CTG ATN WHG TGG TTA GTC GAG CAG AGA C*T*A* G 3'

AG Barcode 8

5' G*A*T* GCA CGC CTT ACG ACT CAT CTN WNH TGA TCT ATN WHG TGG TTA GTC GAG CAG AGA C*T*A* G 3'

AG Barcode 9

5' G*A*T* GCA CGC CTT ACG ACT CAT CTN WNH ATG AGA TGN WHG TGG TTA GTC GAG CAG AGA C*T*A* G 3'

AG Barcode 10

5' G*A*T* GCA CGC CTT ACG ACT CAT CTN WNH GCG AAT TCN WHG TGG TTA GTC GAG CAG AGA C*T*A* G 3'

AG Barcode 11

5' G*A*T* GCA CGC CTT ACG ACT CAT CTN WNH GAT TCC GGN WHG TGG TTA GTC GAG CAG AGA C*T*A* G 3'

AG Barcode 12

5' G*A*T* GCA CGC CTT ACG ACT CAT CTN WNH ATA ATA TAN WHG TGG TTA GTC GAG CAG AGA C*T*A* G 3'

AG Barcode 13

5' G*A*T* GCA CGC CTT ACG ACT CAT CTN WNH AGC ATG CGN WHG TGG TTA GTC GAG CAG AGA C*T*A* G 3'

AG Barcode 14

5' G*A*T* GCA CGC CTT ACG ACT CAT CTN WNH GAT TCA ACN WHG TGG TTA GTC GAG CAG AGA C*T*A* G 3'

AG Barcode 15

5' G*A*T* GCA CGC CTT ACG ACT CAT CTN WNH TAC CTG CTN WHG TGG TTA GTC GAG CAG AGA C*T*A* G 3'

AG Barcode 16

5' G*A*T* GCA CGC CTT ACG ACT CAT CTN WNH GCT AAT CGN WHG TGG TTA GTC GAG CAG AGA C*T*A* G 3'

AG Barcode 17

5' G*A*T* GCA CGC CTT ACG ACT CAT CTN WNH CTC CTT CGN WHG TGG TTA GTC GAG CAG AGA C*T*A* G 3'

AG Barcode 18

5' G*A*T* GCA CGC CTT ACG ACT CAT CTN WNH ACG CTA GCN WHG TGG TTA GTC GAG CAG AGA C*T*A* G 3'

AG Barcode 19

5' G*A*T* GCA CGC CTT ACG ACT CAT CTN WNH GCA GGA CTN WHG TGG TTA GTC GAG CAG AGA C*T*A* G 3'

AG Barcode 20

5' G*A*T* GCA CGC CTT ACG ACT CAT CTN WNH ATT GCT CTN WHG TGG TTA GTC GAG CAG AGA C*T*A* G 3'

AG Barcode 21

5' G*A*T* GCA CGC CTT ACG ACT CAT CTN WNH TAC GCT CGN WHG TGG TTA GTC GAG CAG AGA C*T*A* G 3'

AG Barcode 22

5' G*A*T* GCA CGC CTT ACG ACT CAT CTN WNH ACG CTC CAN WHG TGG TTA GTC GAG CAG AGA C*T*A* G 3'

AG Barcode 23

5' G*A*T* GCA CGC CTT ACG ACT CAT CTN WNH CGG TCA ATN WHG TGG TTA GTC GAG CAG AGA C*T*A* G 3'

AG Barcode 24

5' G*A*T* GCA CGC CTT ACG ACT CAT CTN WNH CGC CTA TTN WHG TGG TTA GTC GAG CAG AGA C*T*A* G 3'

AG Barcode 25

5' G*A*T* GCA CGC CTT ACG ACT CAT CTN WNH TTG CGT TGN WHG TGG TTA GTC GAG CAG AGA C*T*A* G 3'

AG Barcode 26

5' G*A*T* GCA CGC CTT ACG ACT CAT CTN WNH TCC TAA GAN WHG TGG TTA GTC GAG CAG AGA C*T*A* G 3'

AG Barcode 27

5' G*A*T* GCA CGC CTT ACG ACT CAT CTN WNH CAA GAA GGN WHG TGG TTA GTC GAG CAG AGA C*T*A* G 3'

AG Barcode 28

5' G*A*T* GCA CGC CTT ACG ACT CAT CTN WNH TAG AAT TAN WHG TGG TTA GTC GAG CAG AGA C*T*A* G 3'

AG Barcode 29

5' G*A*T* GCA CGC CTT ACG ACT CAT CTN WNH GGC GCC AAN WHG TGG TTA GTC GAG CAG AGA C*T*A* G 3'

AG Barcode 30

5' G*A*T* GCA CGC CTT ACG ACT CAT CTN WNH TAG ATC CGN WHG TGG TTA GTC GAG CAG AGA C*T*A* G 3'

AG Barcode 31

5' G*A*T* GCA CGC CTT ACG ACT CAT CTN WNH CGA GCA GCN WHG TGG TTA GTC GAG CAG AGA C*T*A* G 3'

AG Barcode 32

5' G*A*T* GCA CGC CTT ACG ACT CAT CTN WNH TAA GAT GAN WHG TGG TTA GTC GAG CAG AGA C*T*A* G 3'

AG Barcode 33

5' G*A*T* GCA CGC CTT ACG ACT CAT CTN WNH AGC TCG GAN WHG TGG TTA GTC GAG CAG AGA C*T*A* G 3'

AG Barcode 34

5' G*A*T* GCA CGC CTT ACG ACT CAT CTN WNH TAA CCG AAN WHG TGG TTA GTC GAG CAG AGA C*T*A* G 3'

AG Barcode 35

5' G*A*T* GCA CGC CTT ACG ACT CAT CTN WNH TAT ATC TAN WHG TGG TTA GTC GAG CAG AGA C*T*A* G 3'

AG Barcode 36

5' G*A*T* GCA CGC CTT ACG ACT CAT CTN WNH AAG AGG ATN WHG TGG TTA GTC GAG CAG AGA C*T*A* G 3'

AG Barcode 37

5' G*A*T* GCA CGC CTT ACG ACT CAT CTN WNH ACG TCG AAN WHG TGG TTA GTC GAG CAG AGA C*T*A* G 3'

AG Barcode 38

5' G*A*T* GCA CGC CTT ACG ACT CAT CTN WNH CAT CAT TAN WHG TGG TTA GTC GAG CAG AGA C*T*A* G 3'

AG Barcode 39

5' G*A*T* GCA CGC CTT ACG ACT CAT CTN WNH TTG CAA CTN WHG TGG TTA GTC GAG CAG AGA C*T*A* G 3'

AG Barcode 40

5' G*A*T* GCA CGC CTT ACG ACT CAT CTN WNH TCT AAC TGN WHG TGG TTA GTC GAG CAG AGA C*T*A* G 3'

AG Barcode 41

5' G*A*T* GCA CGC CTT ACG ACT CAT CTN WNH TAT GCC TTN WHG TGG TTA GTC GAG CAG AGA C*T*A* G 3'

AG Barcode 42

5' G*A*T* GCA CGC CTT ACG ACT CAT CTN WNH GTA ATT GCN WHG TGG TTA GTC GAG CAG AGA C*T*A* G 3'

AG Barcode 43

5' G*A*T* GCA CGC CTT ACG ACT CAT CTN WNH GTC TCC GTN WHG TGG TTA GTC GAG CAG AGA C*T*A* G 3'

AG Barcode 44

5' G*A*T* GCA CGC CTT ACG ACT CAT CTN WNH TGC ATG GTN WHG TGG TTA GTC GAG CAG AGA C*T*A* G 3'

AG Barcode 45

5' G*A*T* GCA CGC CTT ACG ACT CAT CTN WNH AGT CCG GTN WHG TGG TTA GTC GAG CAG AGA C*T*A* G 3'

AG Barcode 46

5' G*A*T* GCA CGC CTT ACG ACT CAT CTN WNH TCC TGA TGN WHG TGG TTA GTC GAG CAG AGA C*T*A* G 3'

AG Barcode 47

5' G*A*T* GCA CGC CTT ACG ACT CAT CTN WNH ATC GTC TAN WHG TGG TTA GTC GAG CAG AGA C*T*A* G 3'

AG Barcode 48

5' G*A*T* GCA CGC CTT ACG ACT CAT CTN WNH GGA CGT CCN WHG TGG TTA GTC GAG CAG AGA C*T*A* G 3'

AG Barcode 49

5' G*A*T* GCA CGC CTT ACG ACT CAT CTN WNH CTA CGA GGN WHG TGG TTA GTC GAG CAG AGA C*T*A* G 3'

AG Barcode 50

5' G*A*T* GCA CGC CTT ACG ACT CAT CTN WNH CAA TCC GTN WHG TGG TTA GTC GAG CAG AGA C*T*A* G 3'

**Supplementary Table 3:** Sequences of primers used for barcode amplification

Primer_A_FWD_S502

5' AAT GAT ACG GCG ACC ACC GAG ATC TAC ACC TCT CTA TTC GTC GGC AGC GTC AGA TGT GTA TAA GAG ACA GGC ACG CCT TAC GAC TCA TCT 3'

Primer_A_FWD_S503

5' AAT GAT ACG GCG ACC ACC GAG ATC TAC ACT ATC CTC TTC GTC GGC AGC GTC AGA TGT GTA TAA GAG ACA GGC ACG CCT TAC GAC TCA TCT 3'

Primer A_FWD_S505

5' AAT GAT ACG GCG ACC ACC GAG ATC TAC ACG TAA GGA GTC GTC GGC AGC GTC AGA TGT GTA TAA GAG ACA GGC ACG CCT TAC GAC TCA TCT 3'

Primer A_FWD_S506

5' AAT GAT ACG GCG ACC ACC GAG ATC TAC ACA CTG CAT ATC GTC GGC AGC GTC AGA TGT GTA TAA GAG ACA GGC ACG CCT TAC GAC TCA TCT 3'

Primer A_FWD_S507

5' AAT GAT ACG GCG ACC ACC GAG ATC TAC ACA AGG AGT ATC GTC GGC AGC GTC AGA TGT GTA TAA GAG ACA GGC ACG CCT TAC GAC TCA TCT 3'

Primer A_FWD_S508

5' AAT GAT ACG GCG ACC ACC GAG ATC TAC ACC TAA GCC TTC GTC GGC AGC GTC AGA TGT GTA TAA GAG ACA GGC ACG CCT TAC GAC TCA TCT 3'

Primer A_FWD_S510

5' AAT GAT ACG GCG ACC ACC GAG ATC TAC ACC GTC TAA TTC GTC GGC AGC GTC AGA TGT GTA TAA GAG ACA GGC ACG CCT TAC GAC TCA TCT 3'

Primer A_FWD_S511

5' AAT GAT ACG GCG ACC ACC GAG ATC TAC ACT CTC TCC GTC GTC GGC AGC GTC AGA TGT GTA TAA GAG ACA GGC ACG CCT TAC GAC TCA TCT 3'

Primer A_FWD_S513

5' AAT GAT ACG GCG ACC ACC GAG ATC TAC ACT CGA CTA GTC GTC GGC AGC GTC AGA TGT GTA TAA GAG ACA GGC ACG CCT TAC GAC TCA TCT 3'

1Primer A_FWD_S515

5' AAT GAT ACG GCG ACC ACC GAG ATC TAC ACT TCT AGC TTC GTC GGC AGC GTC AGA TGT GTA TAA GAG ACA GGC ACG CCT TAC GAC TCA TCT 3'

Primer A_FWD_S516

5' AAT GAT ACG GCG ACC ACC GAG ATC TAC ACC CTA GAG TTC GTC GGC AGC GTC AGA TGT GTA TAA GAG ACA GGC ACG CCT TAC GAC TCA TCT 3'

Primer A_FWD_S517

5' AAT GAT ACG GCG ACC ACC GAG ATC TAC ACG CGT AAG ATC GTC GGC AGC GTC AGA TGT GTA TAA GAG ACA GGC ACG CCT TAC GAC TCA TCT 3'

Primer A_FWD_S518

5' AAT GAT ACG GCG ACC ACC GAG ATC TAC ACC TAT TAA GTC GTC GGC AGC GTC AGA TGT GTA TAA GAG ACA GGC ACG CCT TAC GAC TCA TCT 3'

Primer A_FWD_S520

5' AAT GAT ACG GCG ACC ACC GAG ATC TAC ACA AGG CTA TTC GTC GGC AGC GTC AGA TGT GTA TAA GAG ACA GGC ACG CCT TAC GAC TCA TCT 3'

Primer A_FWD_S521

5' AAT GAT ACG GCG ACC ACC GAG ATC TAC ACG AGG CTA TTC GTC GGC AGC GTC AGA TGT GTA TAA GAG ACA GGC ACG CCT TAC GAC TCA TCT 3'

Primer A_FWD_S522

5' AAT GAT ACG GCG ACC ACC GAG ATC TAC ACT TAT GCG ATC GTC GGC AGC GTC AGA TGT GTA TAA GAG ACA GGC ACG CCT TAC GAC TCA TCT 3'

Primer G_RVS_N701

5' CAA GCA GAA GAC GGC ATA CGA GAT TCG CCT TAG TCT CGT GGG CTC GGA GAT GTG TAT AAG AGA CAG GTC TCT GCT CGA CTA ACC AC 3'

Primer G_RVS_N702

5' CAA GCA GAA GAC GGC ATA CGA GAT CTA GTA CGG TCT CGT GGG CTC GGA GAT GTG TAT AAG AGA CAG GTC TCT GCT CGA CTA ACC AC 3'

Primer G_RVS_N703

5' CAA GCA GAA GAC GGC ATA CGA GAT TTC TGC CTG TCT CGT GGG CTC GGA GAT GTG TAT AAG AGA CAG GTC TCT GCT CGA CTA ACC AC 3'

Primer G_RVS_N704

5' CAA GCA GAA GAC GGC ATA CGA GAT GCT CAG GAG TCT CGT GGG CTC GGA GAT GTG TAT AAG AGA CAG GTC TCT GCT CGA CTA ACC AC 3'

Primer G_RVS_N705

5' CAA GCA GAA GAC GGC ATA CGA GAT AGG AGT CCG TCT CGT GGG CTC GGA GAT GTG TAT AAG AGA CAG GTC TCT GCT CGA CTA ACC AC 3'

Primer G_RVS_N706

5' CAA GCA GAA GAC GGC ATA CGA GAT CAT GCC TAG TCT CGT GGG CTC GGA GAT GTG TAT AAG AGA CAG GTC TCT GCT CGA CTA ACC AC 3'

Primer G_RVS_N707

5' CAA GCA GAA GAC GGC ATA CGA GAT GTA GAG AGG TCT CGT GGG CTC GGA GAT GTG TAT AAG AGA CAG GTC TCT GCT CGA CTA ACC AC 3'

Primer G_RVS_N710

5' CAA GCA GAA GAC GGC ATA CGA GAT CAG CCT CGG TCT CGT GGG CTC GGA GAT GTG TAT AAG AGA CAG GTC TCT GCT CGA CTA ACC AC 3'

Primer G_RVS_N711

5' CAA GCA GAA GAC GGC ATA CGA GAT TGC CTC TTG TCT CGT GGG CTC GGA GAT GTG TAT AAG AGA CAG GTC TCT GCT CGA CTA ACC AC 3'

Primer G_RVS_N712

5' CAA GCA GAA GAC GGC ATA CGA GAT TCC TCT ACG TCT CGT GGG CTC GGA GAT GTG TAT AAG AGA CAG GTC TCT GCT CGA CTA ACC AC 3'

Primer G_RVS_N714

5' CAA GCA GAA GAC GGC ATA CGA GAT TCA TGA GC G TCT CGT GGG CTC GGA GAT GTG TAT AAG AGA CAG GTC TCT GCT CGA CTA ACC AC 3'

Primer G_RVS_N715

5' CAA GCA GAA GAC GGC ATA CGA GAT CCT GAG ATG TCT CGT GGG CTC GGA GAT GTG TAT AAG AGA CAG GTC TCT GCT CGA CTA ACC AC 3'

Primer G_RVS_N716

5' CAA GCA GAA GAC GGC ATA CGA GAT TAG CGA GTG TCT CGT GGG CTC GGA GAT GTG TAT AAG AGA CAG GTC TCT GCT CGA CTA ACC AC 3'

Primer G_RVS_N718

5' CAA GCA GAA GAC GGC ATA CGA GAT GTA GCT CCG TCT CGT GGG CTC GGA GAT GTG TAT AAG AGA CAG GTC TCT GCT CGA CTA ACC AC 3'

Primer G_RVS_N719

5' CAA GCA GAA GAC GGC ATA CGA GAT TAC TAC GCG TCT CGT GGG CTC GGA GAT GTG TAT AAG AGA CAG GTC TCT GCT CGA CTA ACC AC 3'

Primer G_RVS_N720

5' CAA GCA GAA GAC GGC ATA CGA GAT AGG CTC CGG TCT CGT GGG CTC GGA GAT GTG TAT AAG AGA CAG GTC TCT GCT CGA CTA ACC AC 3'
